# Recent odor experience selectively modulates olfactory sensitivity across the glomerular output in the mouse olfactory bulb

**DOI:** 10.1101/2024.07.21.604478

**Authors:** Narayan Subramanian, Lee Min Leong, Paria Salemi Mokri Boukani, Douglas A. Storace

## Abstract

Although animals can reliably locate and recognize odorants embedded in complex environments, the neural circuits for accomplishing these tasks remain incompletely understood. Adaptation is likely to be important as it could allow neurons in a brain area to adjust to the broader sensory environment. Adaptive processes must be flexible enough to allow the brain to make dynamic adjustments, while maintaining sufficient stability so that organisms do not forget important olfactory associations. Processing within the mouse olfactory bulb is likely involved in generating adaptation, although there are conflicting models of how it transforms the glomerular output of the mouse olfactory bulb. Here we performed 2-photon Ca^2+^ imaging from mitral/tufted glomeruli in awake mice to determine the time course of recovery from adaptation, and whether it acts broadly or selectively across the glomerular population. Individual glomerular responses, as well as the overall population odor representation was similar across imaging sessions. However, odor-concentration pairings presented with interstimulus intervals upwards of 30-s evoked heterogeneous adaptation that was concentration-dependent. We demonstrate that this form of adaptation is unrelated to variations in respiration, and olfactory receptor neuron glomerular measurements indicate that it is unlikely to be inherited from the periphery. Our results indicate that the olfactory bulb output can reliably transmit stable odor representations, but recent odor experiences can selectively shape neural responsiveness for upwards of 30 seconds. We propose that neural circuits that allow for non-uniform adaptation across mitral/tufted glomerular could be important for making dynamic adjustments in complex odor environments.

## Introduction

Although animals can reliably locate and recognize odorants embedded in complex chemical contexts, the neural circuits for accomplishing these tasks remain incompletely understood (Gross-Isseroff and Lancet, 1988; Uchida and Mainen, 2007; Homma et al., 2009). The ability to attend to, and segment different kinds of odorant information would be facilitated by adaptive processes within the brain that could adjust responsiveness based on an organism’s sensory environment and internal state (Verhagen et al., 2007; Wark et al., 2007; Whitmire and Stanley, 2016; Weber and Fairhall, 2019; Benda, 2021). Such adaptive processes should be flexible enough to allow the brain to make dynamic adjustments to different sensory environments, while maintaining sufficient stability such that organisms do not forget important olfactory associations. The brain areas and the mechanisms underlying how adaptation transforms olfactory representations remain incompletely understood (Gottfried, 2010; Martelli and Storace, 2021).

Odors evoke varying degrees of neural activity across the olfactory receptor neuron population, yielding a combinatorial code that is transmitted to the olfactory bulb (Duchamp-Viret et al., 1999; Wachowiak and Cohen, 2001; Fried et al., 2002; Storace and Cohen, 2017; Zak et al., 2020). Each olfactory receptor type typically maps to one or two regions of neuropil called glomeruli in the olfactory bulb (Mombaerts et al., 1996; Potter et al., 2001). Mitral and tufted cells each innervate a single glomerulus where they receive olfactory receptor input, and send a transformed sensory representation to the rest of the brain (Igarashi et al., 2012; Storace and Cohen, 2017; Storace et al., 2019; Storace and Cohen, 2021).

Although adaptation occurs at the level of olfactory receptor neurons, adaptation is present in mitral cells that is not necessarily inherited from the periphery (Torre et al., 1995; Kurahashi and Menini, 1997; Dietz and Murthy, 2005; Chaudhury et al., 2010; Storace and Cohen, 2021). There are conflicting models of how adaptation impacts the transmission of olfactory sensory information from the bulb to the rest of the brain. There currently exists data to support the presence of adaptation that occurs gradually over days and broadly impacts the output of the bulb, as well as adaptation that is shorter-lasting and selective in which glomerular output channels are affected (Kato et al., 2012; Ogg et al., 2015; Ogg et al., 2018; Storace and Cohen, 2021).

We tested these models using *in vivo* 2-photon Ca^2+^ imaging from the apical dendrites of mitral/tufted cells innervating the olfactory bulb glomerular layer in awake mice. A panel of odors at different concentrations were delivered at different interstimulus intervals on the same day, or during imaging sessions performed on different days. Odor representations across the mitral/tufted glomerular population had similar amplitudes and were well correlated across trials recorded on the same day separated by a minimum of 3 minutes, and in trials measured on different imaging days. However, adaptation was present in response to odors delivered on shorter timescales that was selective across the mitral/tufted glomerular population. Within the same imaging field of view, some glomeruli responded similarly to each odor presentation, while others changed significantly. This adaptation was strongest when odors were presented with a 6-s interstimulus interval and concentration-dependent. Extending the interstimulus interval to 30-s resulted in an incomplete recovery from adaptation. Measurements of animal respiration and odor-evoked activity in olfactory receptor neuron glomeruli suggest that adaptation is unlikely to reflect variations in the organismal state or sensory input. Our results indicate that the mouse olfactory bulb transmits reliable representations of olfactory stimuli across time, although neural processes can selectively mediate sensitivity adjustments based on recent odor exposure for upwards of 30 seconds. We propose that dynamic adaptation in subsets of glomeruli would be useful for making dynamic adjustments to complex odor environments (Gottfried, 2010; Martelli and Storace, 2021).

## Results

### Odors evoke similar responses in mitral/tufted glomeruli when measured during the same imaging session or across different days

To selectively image from mitral/tufted glomeruli, the Tbx21-Cre transgenic mouse line (that expresses cre recombinase in mitral/tufted cells) was mated to the Ai148 cre-dependent GCaMP6f reporter line (Mitsui et al., 2011; Daigle et al., 2018; Koldaeva et al., 2021; Storace and Cohen, 2021). The resulting transgenic offspring were histologically confirmed to express GCaMP6f in mitral/tufted cells and their corresponding glomerular tufts (**Fig. 1A**). During *in vivo* imaging in awake mice, mitral/tufted glomeruli were clearly visible in the mean fluorescence which allowed for routine tracking of the same glomeruli across multiple experimental sessions (**Fig. 1B**, the top row is the mean fluorescence from the same field of view on different days). Frame subtraction images were generated by subtracting the average of the frames during the odor stimulus from the average of the frames prior to the odor. Different odors evoked different patterns of activity across the glomerular population while the same odors evoked activity patterns that were comparable in amplitude and spatial arrangement on different imaging sessions (**Fig. 1B**, bottom row, compare methyl valerate on days 1 6 and 21, isoamyl acetate on days 8 and 9).

**Figure 1:**
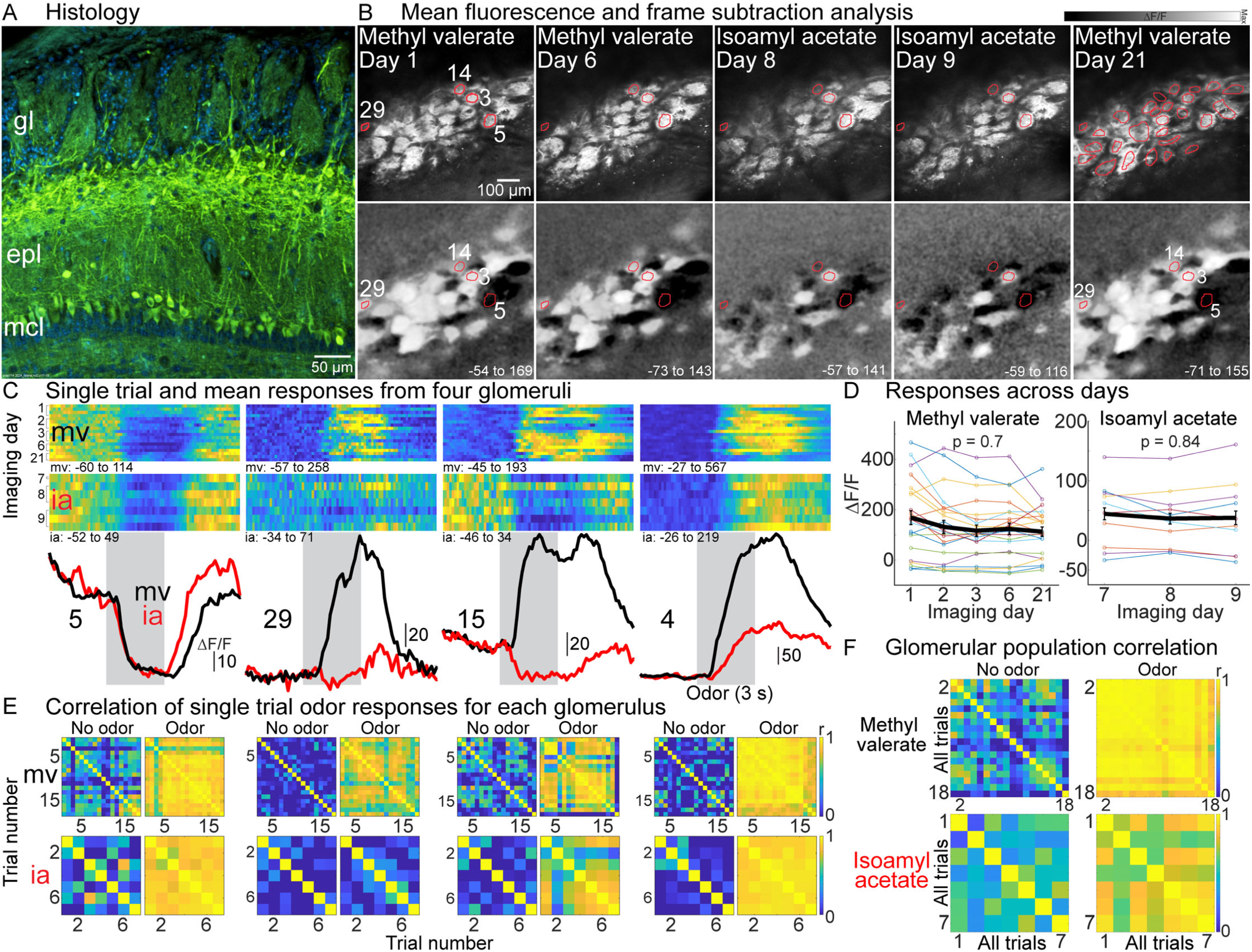
(A) Histology illustrating GCaMP6f expression in mitral/tufted cells. (B) Mean fluorescence (top) and frame subtraction images (bottom) from different days. The ΔF/F scaling is fixed for each odor and the ΔF/F range is in the bottom right of each panel. (C) Single trial and mean response from four glomeruli to two odors presented at 3.5% of saturated vapor. The heat map intensity scaling fixed for all trials for each glomerulus. The ΔF/F range for each heat map is beneath each panel. (D) Odor responses from individual glomeruli (thin lines) and the population mean (thick line) on each imaging day. (E) Correlation of signals before and during odor stimulation. (F) Correlation of the glomerular population response for each pair of imaging trials. gl, glomerular layer; epl, external plexiform layer; mcl, mitral cell layer.

Fluorescence time course measurements from trials separated by at least 3 minutes during the same recording session, and trials carried out on different imaging days had similar time course dynamics and response amplitudes (**Fig. 1C**, the single trial heat maps are scaled to the maximum ΔF/F response for each glomerulus-odor pairing). Although individual glomeruli sometimes exhibited variability on different days (e.g., **Fig. 1C**, ROI 14), the mean response across all glomeruli within a field of view was not significantly different for a given odor across different imaging sessions (**Fig. 1D**, the thin lines are measurements from individual glomeruli, methyl valerate p = 0.7; isoamyl acetate p = 0.84, the analysis only includes glomeruli in the field of view that responded to odor stimulation with at least a 3 standard deviation change from baseline).

In glomeruli that responded to the odor, responses were highly correlated during (but not prior to) odor stimulation in trials measured on the same day or on different days (**Fig. 1E**). The mean correlation of individual glomeruli in this preparation in response to methyl valerate before and during odor stimulation was 0.08 ± 0.01 and 0.62 ± 0.02, respectively (range air: −0.02 – 0.34, odor: 0.04-0.95, N = 29 glomeruli). Similar results were obtained from individual glomeruli in response to isoamyl acetate in this preparation (air: 0.11 ± 0.04, range −0.06-0.5; odor: 0.34 ± 0.08, range −0.02 – 0.9). The mean glomerular response during odor stimulation was highly correlated in single trials measured on the same or on different days in comparison with the time preceding the odor stimulus (**Fig. 1F**). The mean correlation between measurements from trials measured on the same day or on different days were not statistically different from one another (**Fig. 1F**, methyl valerate: same day trials r = 0.95 ± 0.01 N = 24; different day trials r = 0.93 ± 0.01, N = 153; p = 0.06; isoamyl acetate, same day trials r = 0.39 ± 0.1 N = 7; different day r = 0.43 ± 0.05, N = 15; p = 0.9).

In each preparation-odor pairing, imaging was carried out on up to 5 different days (4 ± 0.25 imaging sessions; N = 16), separated by an average of 2.2 ± 0.8 days (range included 1-18 days between imaging sessions within the same preparation-odor pairings). The mean glomerular response was not significantly different on different imaging days for individual preparation-odor pairing (p-values ranged from 0.16-0.97 using a Kruskal-wallis test). The population mean amplitude (ΔF/F) was not significantly different when imaging sessions were binned relatively (**Fig. 2A**, p = 0.66), or based on the absolute number of days between imaging sessions (**Fig. 2B**, p = 0.8), nor in a separate analysis restricted to preparations imaged on two consecutive days (**Fig. 2B**, comparison between data points on days 1-2, p = 0.38; 10/16 preparation-odor pairings). For trials measured during the same imaging sessions or on different days, the correlation of the mean glomerular response was not significantly different from one another (**Fig. 2C**, same day: 0.72 ± 0.05, different day: 0.72 ± 0.05, p = 0.66).

**Figure 2:**
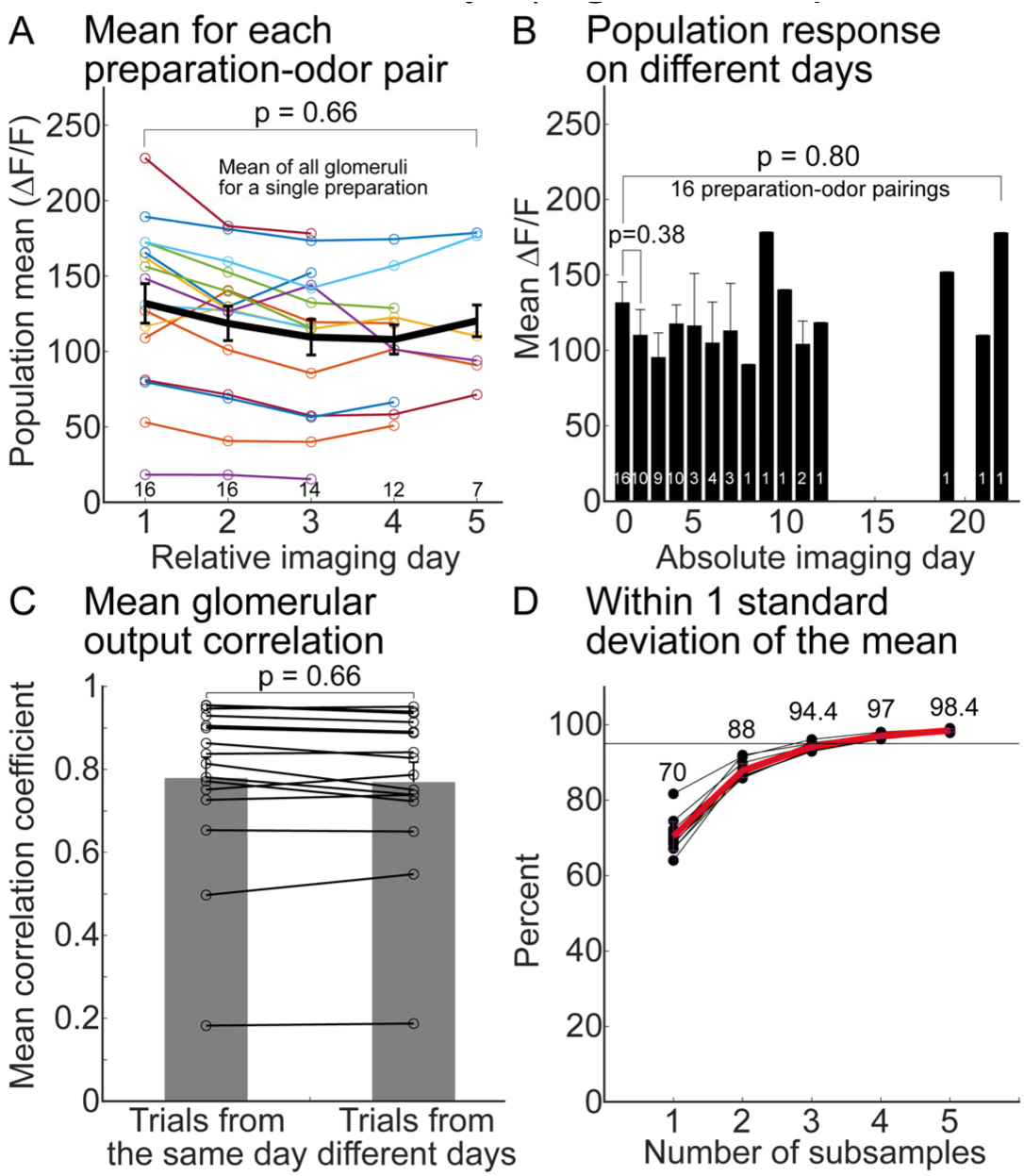
(A) Mean response across all responsive glomeruli for individual preparation-odor pairings (thin lines) and the overall mean (thick line) for different imaging sessions. (B) The results from panel A binned as a function of the number of days since the 1st imaging session. The number of preparation-odor pairings included in each mean is indicated at the bottom of A-B. (C) The mean correlation of the population response in each preparation-odor pairing for trials recorded on the same (left bar) or different days (right bar). (D) The percentage of 1000 subsamples that came within 1 standard deviation of the unsampled mean.

Although these data demonstrate that the responses of individual mitral/tufted glomeruli as well as the population mean are similar in amplitude and highly correlated during odor stimulation, there is unexplained variability across trials (e.g., **Fig. 1C**, ROI 29 mv). We estimated the number of trials required to obtain a representative response for a given glomerulus by randomly (with replacement) selecting between 1-6 samples from all trials for each preparation-odor pairing. This process was repeated 1000 times, each time the mean of the subsampled distribution was compared to the unsampled mean. Subsampling at least 4 trials resulted in a mean that was within 1 standard deviation of the unsampled mean in 97 % of the 1000 repetitions (**Fig. 2D**, thin lines are from different preparation-odor pairings). This analysis includes measurements in 11.43 ± 1.76 trials per preparation-odor pairing measured at 3.2% of saturated vapor (range of 4-31 trials across all imaging sessions). This result suggests that 4 repetitions of a stimulus provide sufficient information to characterize the odor response of a single glomerulus.

### Adaptation is present in subsets of MTC glomeruli in response to 6-s interstimulus intervals

2-photon imaging was previously used to demonstrate that mitral/tufted glomeruli exhibit adaptation in response to odors presented with an interstimulus interval of 6-s in anesthetized mice (Storace and Cohen, 2021). Because mitral/tufted cell activity and respiratory patterns can vary substantially in awake and anesthetized states, we tested whether it is present in awake mice (Rinberg et al., 2006; Kato et al., 2012; Wachowiak et al., 2013). The effect of repeated odor presentations across the glomerular population is illustrated using a frame subtraction analysis in preparations in which multiple odors were tested (**Fig. 3**). The average of the frames during the 1^st^ and 3^rd^ odor presentation subtracted from the average of the frames preceding each odor stimulus visualizes glomeruli that are activated and suppressed as white and black, respectively (**Fig. 3B-D**, 1^st^ and 3^rd^ odor). The difference of the average of the frames during the 1^st^ and 3^rd^ odor presentation visualizes non-adapting and adapting glomeruli as gray and black, respectively (**Fig. 3B-D**, Difference). The pattern of glomeruli activated by the odor, and the degree to which they adapted changed in an odor-specific way (**Fig. 3B-D**, compare 1^st^ odor presentation and difference maps for different odors). Similar results were obtained in all preparations in which we measured responses to at least two different odors (N = 6 preparations).

**Figure 3:**
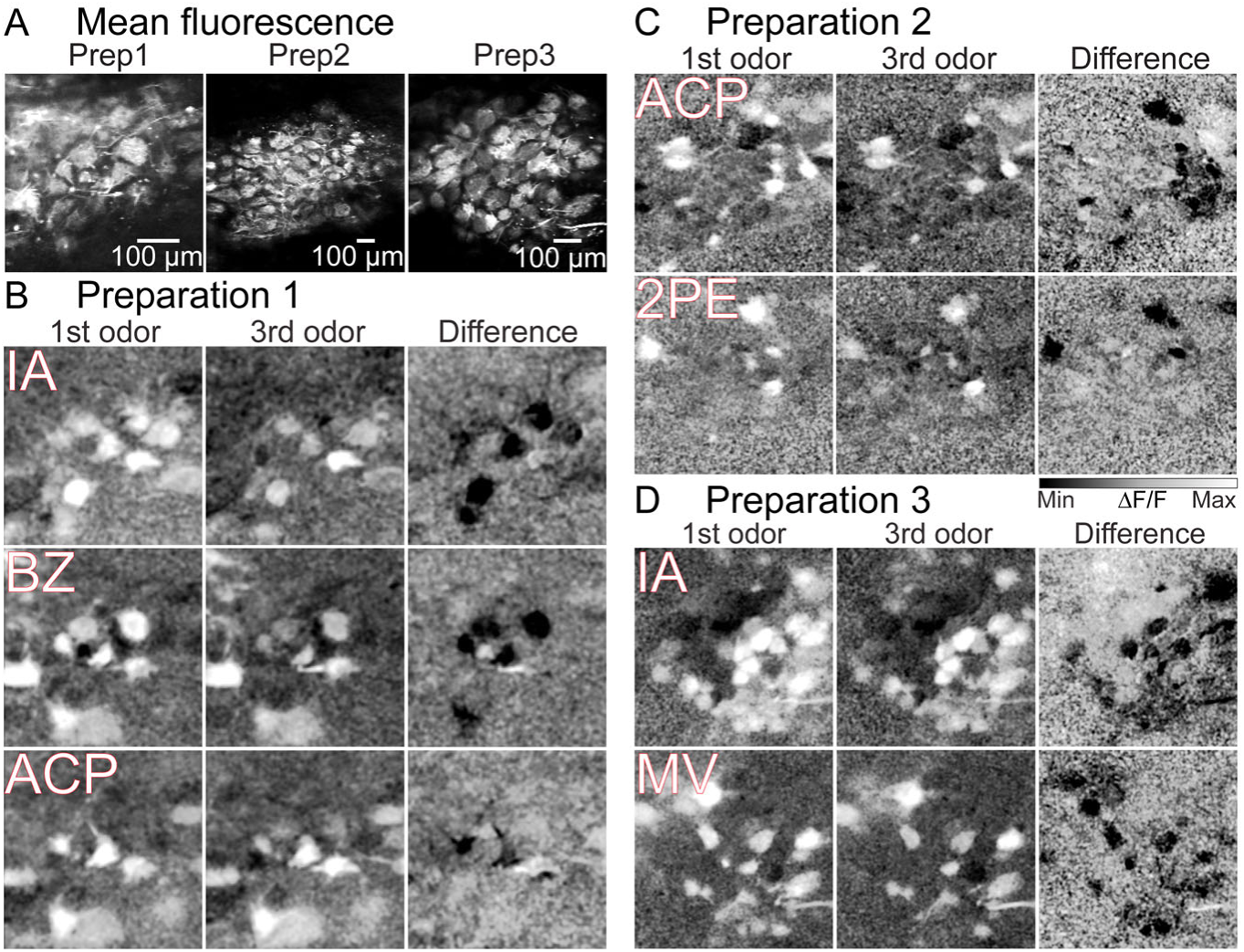
(A) Mitral/tufted glomerular fluorescence in three preparations. (B-D) Frame subtraction images in response to odors presented at 3.5% of saturated vapor for the preparations in panel A. (1st and 3rd) Response to the 1st and 3rd odor presentations. The intensity-scaling is fixed to the same range. (Difference) Difference between the 1st and 3rd presentation. IA, isoamyl acetate; BZ, benzaldehyde; ACP, acetophenone; MV, methyl valerate; 2PE, 2-phenylethanol.

Odor-specific responses and adaptation is further illustrated in single trial and mean fluorescence time course measurements from three preparations (**Fig. 4**). The same odor evoked different non-adapting and adapting responses in glomeruli imaged simultaneously in the same field of view (**Fig. 4A, C, E**). Non-adapting glomeruli included those that were excited or suppressed by the odor (**Fig. 4A** methyl valerate ROI 2; **Fig. 4C** isoamyl acetate ROI 6; **Fig. 4E** methyl valerate ROIs 29 and 1). All preparation-odor pairings had other glomeruli in the same field of view that responded to the 3^rd^ odor presentation differently than to the 1^st^. This included glomeruli which returned to baseline following odor removal (**Fig. 4A**, 2-phenylethanol ROI 63; **Fig. 4C** isoamyl acetate ROI 51), and others that exhibited a slow decaying response (**Fig. 4A**, acetophenone ROI 63). Other adapting glomeruli responded with a calcium increase in response to the 1^st^ presentation, and became suppressed to subsequent presentations (**Fig. 4A**, acetophenone ROI 9; **Fig. 4E**, methyl valerate ROIs 8 and 10).

**Figure 4:**
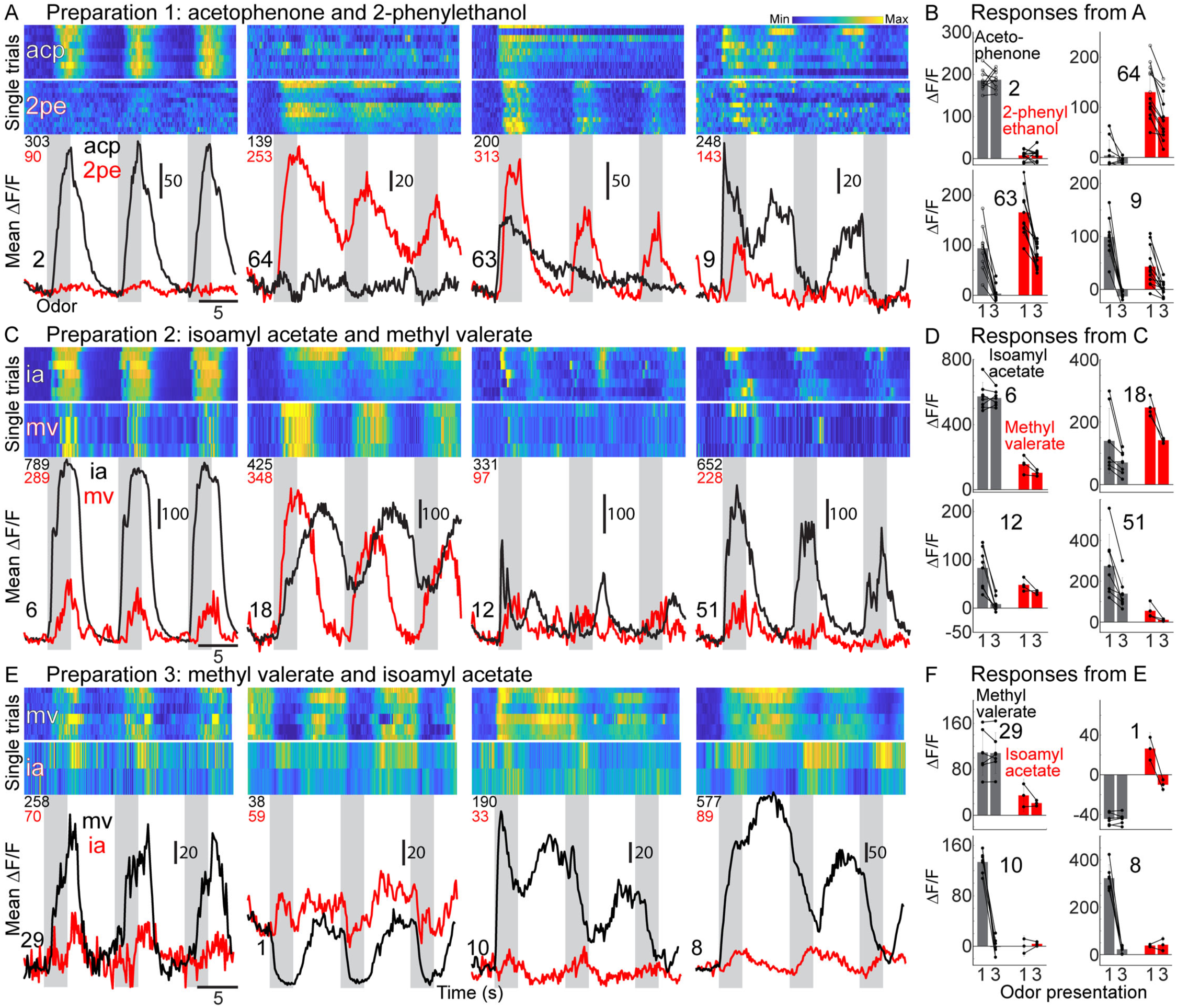
(A, C, E) Single trial and mean fluorescence time course from 4 glomeruli in 3 preparations in response to two odors. All four glomeruli in each preparation were measured simultaneously in the same fields of view. The odor command signal is illustrated with gray bars. The numbers underneath the heat maps indicate the max ΔF/F across all trials for that each odor. (B, D, F) Mean and single trial (connected lines) odor responses from the glomeruli in panels A, C, and E.

A different odor could evoke no response (**Fig. 4A**, ROI 2), a different amplitude response (**Fig. 4C**, ROI 6), and similar or different adapting or non-adapting responses in the same glomerulus (**Fig. 4A**, ROI 63 versus **Fig. 4E**, ROI 8). A quantification of the amplitudes of single trials illustrates that the adapting or non-adapting characteristics of glomeruli are stable across individual trials for a given odor (**Fig. 4B, D, F**).

We tested whether the rate and numbers of odor inhalations could account for mitral/tufted glomerular adaptation. Single trial measurements from glomeruli simultaneously imaged in the same field of view illustrate the presence of non-adapting and adapting glomeruli during similar respiratory patterns during each odor presentation (**Fig. 5A**). Additional representative example recordings of respiration during imaging trials illustrate that respiration activity did not systematically change during our odor stimulation paradigm (**Fig. 5B**). The mean number of inhalations during the 1^st^ and 3^rd^ odor presentation was not significantly different in an exemplar preparation (**Fig. 5C**, 67 trials, p = 0.3; counts from the same trial are connected by a line), across a population of imaging trials (**Fig. 5D**, 1^st^: 10.4 ± 0.5; 3^rd^: 10.2 ± 0.5, p = 0.75, N = 186 trials in 8 preparations), and for comparisons of trials within individual preparations (**Fig. 5E**; p-values range between 0.08-0.62 for individual preparation comparisons). The inter-inhalation interval, which quantifies the time between two subsequent inhalations (Wesson et al., 2008a) was not significantly different in the exemplar preparation (**Fig. 5F**, p = 0.47). Although there was a small increase in the inter-inhalation interval between the 1^st^ and 3^rd^ presentation when collapsing all inhalation pairs in the data set (**Fig. 5G**, P1: 284.7 ± 3.2, N = 1751; P3: 290.5 ± 2.9, N = 1723, p < 0.05), individual preparations comparisons were not significantly different (**Fig. 5H**, p-values range between 0.09-0.84 for individual preparations).

**Figure 5:**
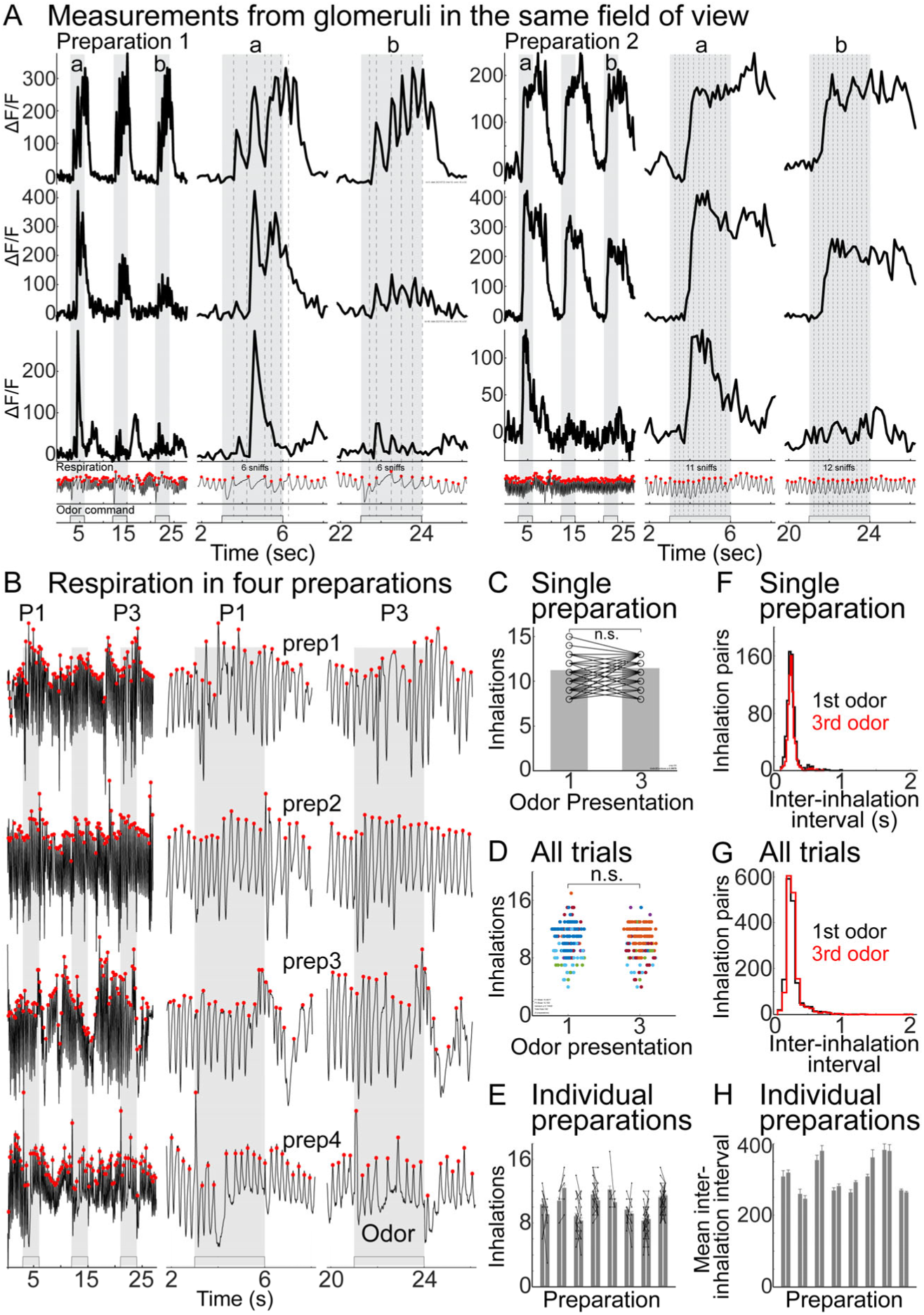
(A) Single trial measurements from glomeruli in two preparations. The odor command timing is indicated by the gray bar. Respiration timing is indicated with red circles and vertical lines indicate respiration during the odor presentation. (B) Respiration in four awake mice. (C) Inhalation counts during odor stimulation in one preparation. (D) Inhalation counts during odor stimulation for all trials. (E) Mean inhalations per trial in 8 preparations. Each line indicates the number of inhalations within a trial. (F) Inter-inhalation interval for the same preparation from panel C. (G) Inter-inhalation interval for all inhalation-pairs in panel D. (H) Mean inter-inhalation interval during the 1st and 3rd odor presentation for 8 preparations.

To confirm that our olfactometer produces repeatable odor stimuli, we used a photoionization detector (PID) to measure the response to methyl valerate presented at different air and liquid dilutions. Odor presentations with a 6-s interstimulus evoked similar time course signals, the amplitudes of which were not significantly different across presentations (**Fig. 6A-B**, p > 0.71 for all comparisons). The PID amplitude measured at different air dilutions (% of saturated vapor) was relatively linear for the undiluted odorant and when it was diluted 1:10 or 1:100 in mineral oil (**Fig. 6C**, *black, red, and blue points*). Interestingly, a 1:10 dilution in mineral oil at 10% of saturated vapor reduced the PID signal by a factor of ∼2, an amplitude near the response evoked by pure odor at 6 % of saturated vapor (**Fig. 6D**). A 1:100 dilution in mineral oil delivered at 10 % of saturated vapor reduced the PID amplitude by a factor of ∼15, an amplitude comparable to the response evoked by pure odor at delivered at 3 % of saturated vapor (**Fig. 6C-D**).

**Figure 6:**
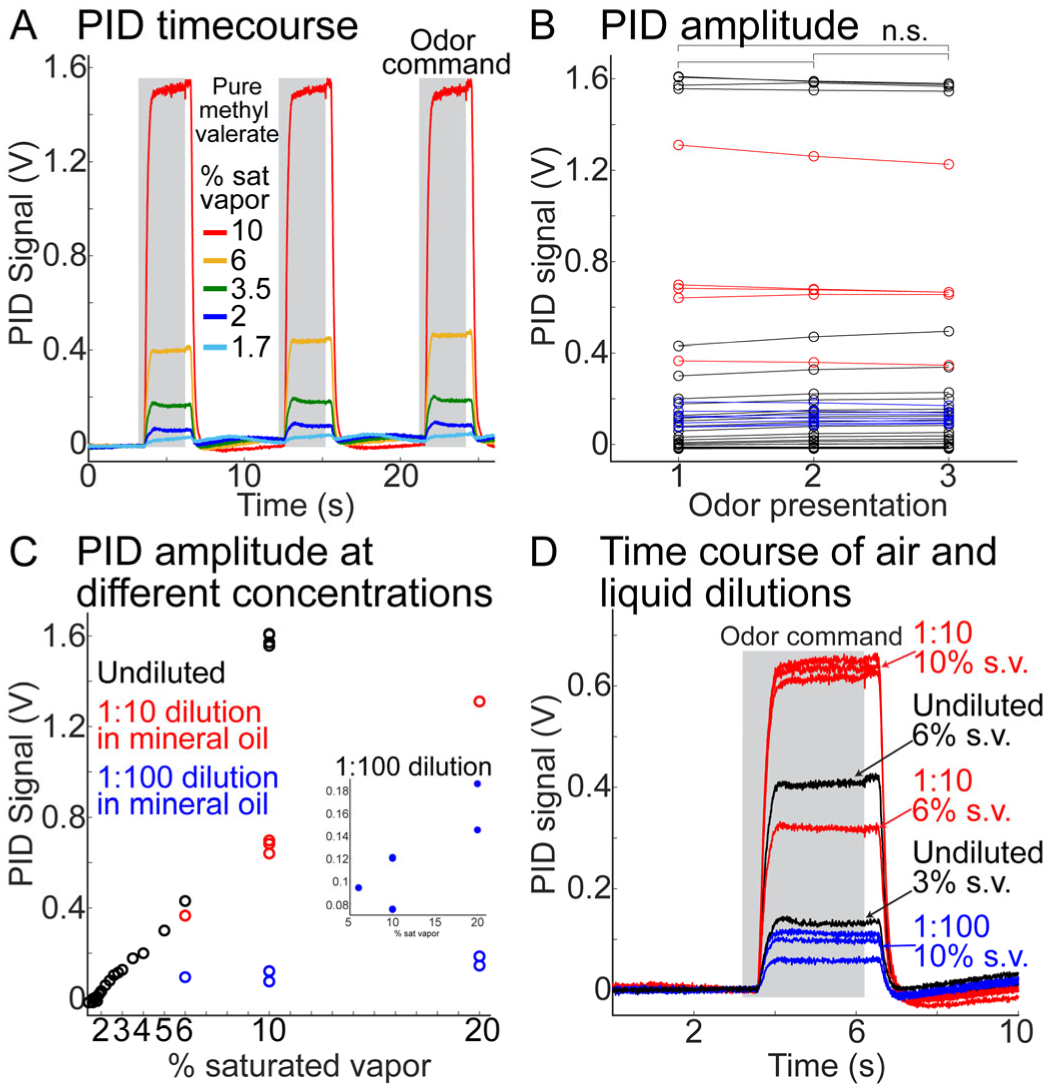
(A-B) PID time course (A) and amplitude response (B) to repeated presentations of pure methyl valerate at different air dilutions. (C) PID amplitude in response to different air and liquid dilutions of methyl valerate. The 1:100 dilution data are expanded in the inset. (D) PID time course to air and liquid dilutions.

### Adaptation is a complicated function of concentration and interstimulus interval

We examined how interstimulus interval and odor concentration influenced the magnitude of adaptation and the degree of recovery. For odors presented at the highest concentration, extending the interstimulus interval from 6-s to 12-s to 30-s evoked similar responses in the non-adapting glomeruli and graded but often incomplete recoveries in adapting glomeruli (**Fig. 7**). The effect of concentration is illustrated on four different glomeruli measured in a different exemplar preparation (**Fig. 8**). Lower concentrations evoked minimal adaptation regardless of the interstimulus interval (**Fig. 8**, *blue traces*). At the 6-s interstimulus interval condition, higher concentrations resulted in non-uniform changes across the glomerular population.

**Figure 7:**
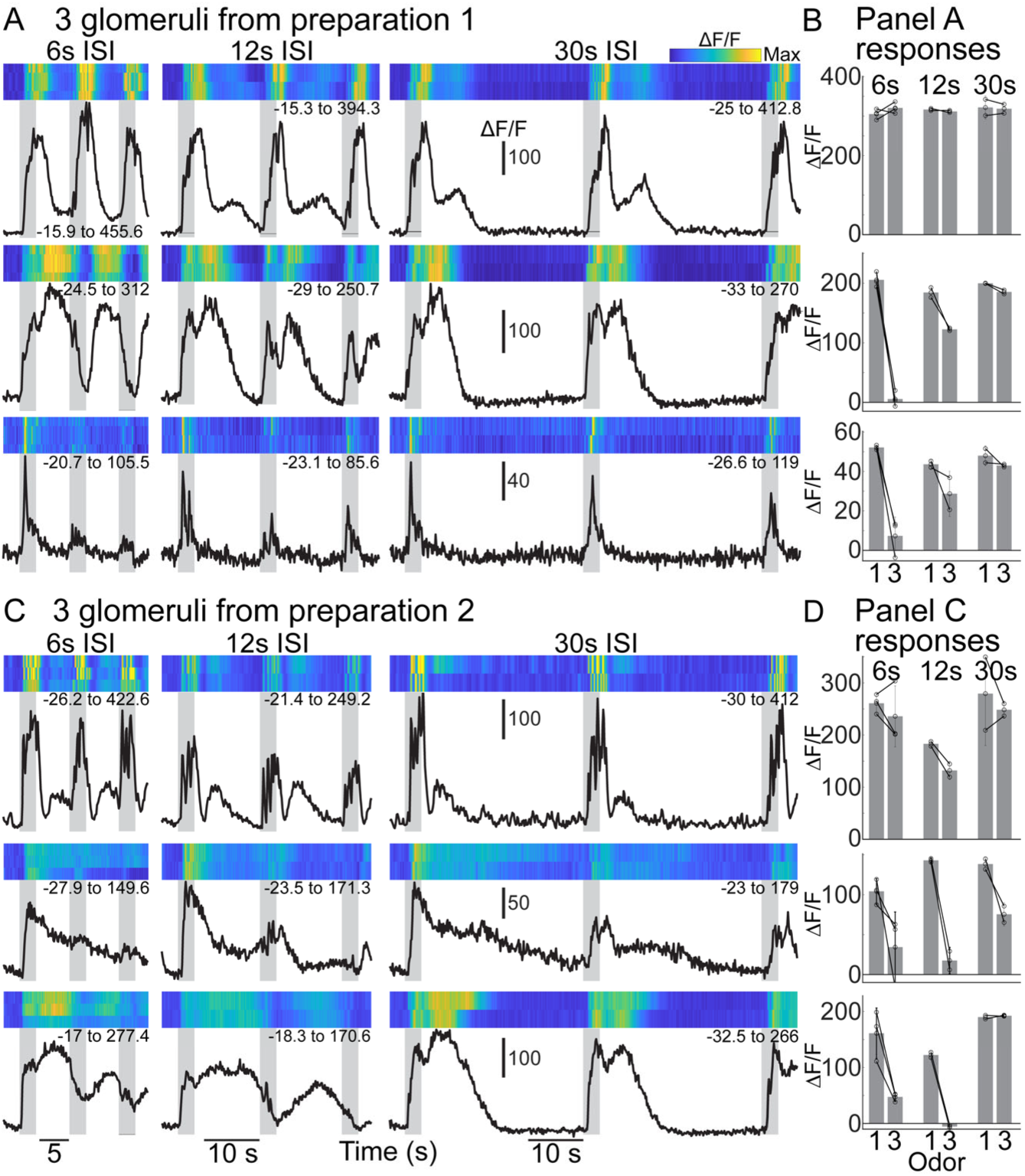
**(A, C)** Single trail and mean odor responses from glomeruli in the same field of view in two preparations. Methyl valerate was presented at 6-s, 12-s and 30-s interstimulus intervals in both preparations. The ΔF/F range for each glomerulus is indicated beneath each heatmap. **(B, D)** Response amplitudes for the measurements in panels **A** and **C**.

**Figure 8:**
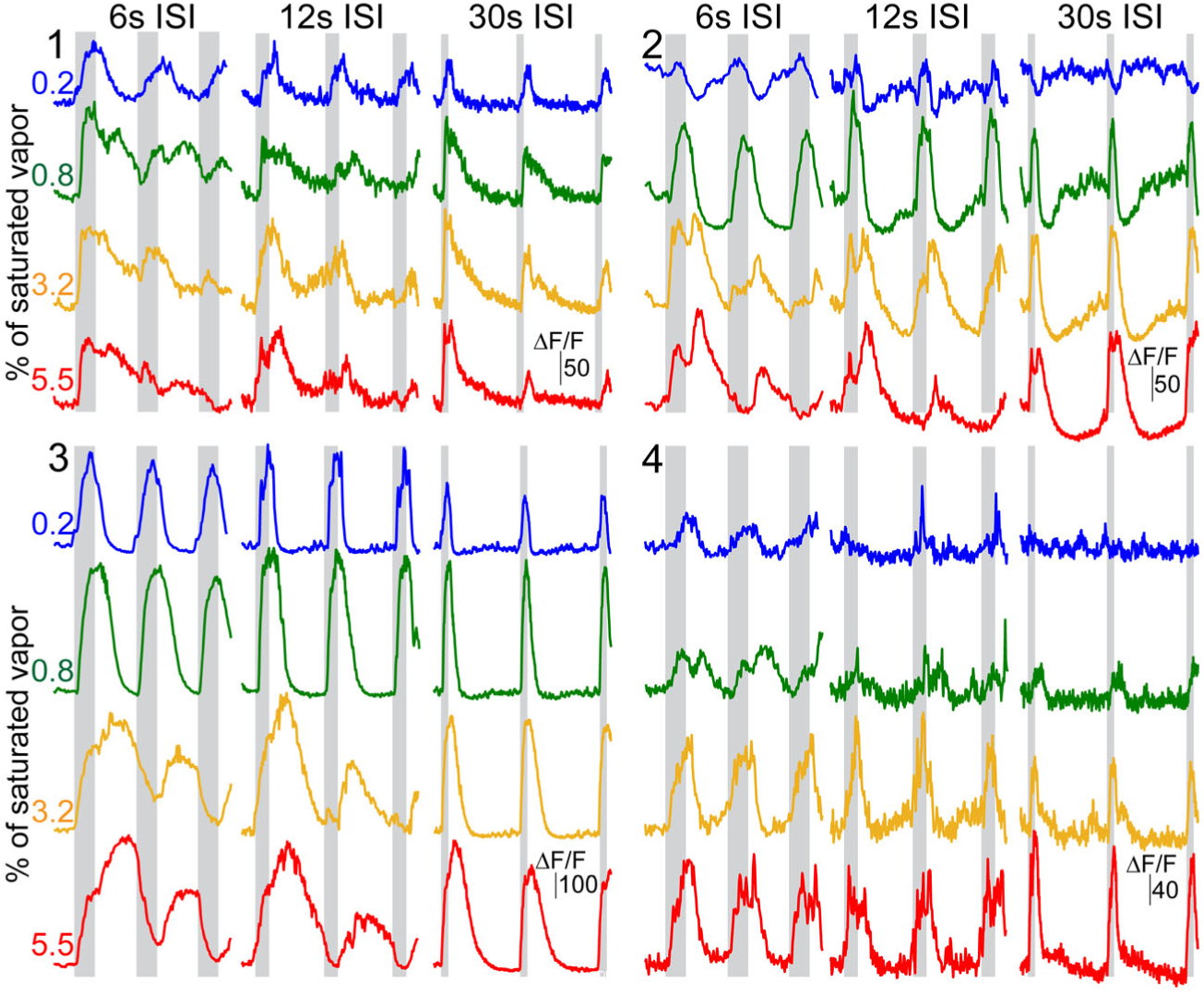
Mean odor responses from four glomeruli recorded simultaneously in the same field of view to methyl valerate presented at different concentrations and interstimulus intervals.

Repeated presentations at 0.8% of saturated vapor evoked progressively weaker responses in glomerulus 1, while glomeruli 2-4 remained stable (**Fig. 8**, green traces). Higher concentrations evoked progressively stronger and more complex adaptation in glomeruli 1-3 (**Fig. 8**, green, orange, and red traces).

At lower concentrations glomerulus 2 dipped below baseline after the stimulus offset, where higher concentrations evoked an excitatory offset response with a slow decay and progressively weaker responses (**Fig. 8**, ROI 2). Glomerulus 3 transitioned from non-adapting to polarity switching at higher concentrations (**Fig. 8**, ROI 3). Glomerulus 4 remained non-adapting at each concentration that evoked a response (e.g., **Fig. 8** ROI 4). The effect of interstimulus interval at different concentrations was consistent with the results in **Fig. 7** in which there was a graded but often incomplete recovery (**Fig. 8**, 12s and 30s ISI).

We quantified the heterogeneity of adaptation in individual glomeruli in preparations in which the response to each concentration was measured in at least 4 trials for both the 6-s and 30-s interstimulus intervals (trials per odor-concentration condition: mean of 7.4 ± 1.3, range of 4-17). This data set includes 358 glomerular measurements in 7 preparation-odor pairings from 6 mice; 29-64 glomeruli per preparation; includes responses to methyl valerate, isoamyl acetate, benzaldehyde, acetophenone and 2-phenylethanol). In each preparation-odor pairing, at least one glomerulus was present that did not adapt, and at least one glomerulus exhibited statistically significant adaptation.

The mean response of individual glomeruli to the 1^st^ and 3^rd^ odor presentation are illustrated in scatterplots for 3 different preparations (**Fig. 9A-C**, glomeruli exhibiting significant adaptation are indicated in red). For each preparation, more individual glomeruli exhibit a smaller and significantly different response to the 3^rd^ odor presentation at the higher concentrations and 6-s interstimulus interval (**Fig. 9A-C**, 6s-ISI). Extending the interstimulus interval to 30-s resulted in a recovery where the response to the 3^rd^ presentation was more like the 1^st^, and there were fewer significant differences (**Fig. 9A-C**, 30-s ISI). Of the 201 glomeruli that responded to odor stimulation, 88 exhibited statistically significant adaptation at some concentration (p < 0.05). The proportion of significantly adapting glomeruli within the same field of view increased with odor concentration for both 6-s and 30-s interstimulus intervals (**Fig. 9D-E**). For odors delivered with a 6-s interstimulus interval, the proportion of non-adapting glomeruli was only significantly larger at the lowest tested concentration (**Fig. 9D**, p-values indicated in graph). In contrast, odors delivered with a 30-s interstimulus evoked a significantly larger proportion of non-adapting glomeruli at all concentrations (**Fig. 9E**, p-values indicated in graph).

**Figure 9:**
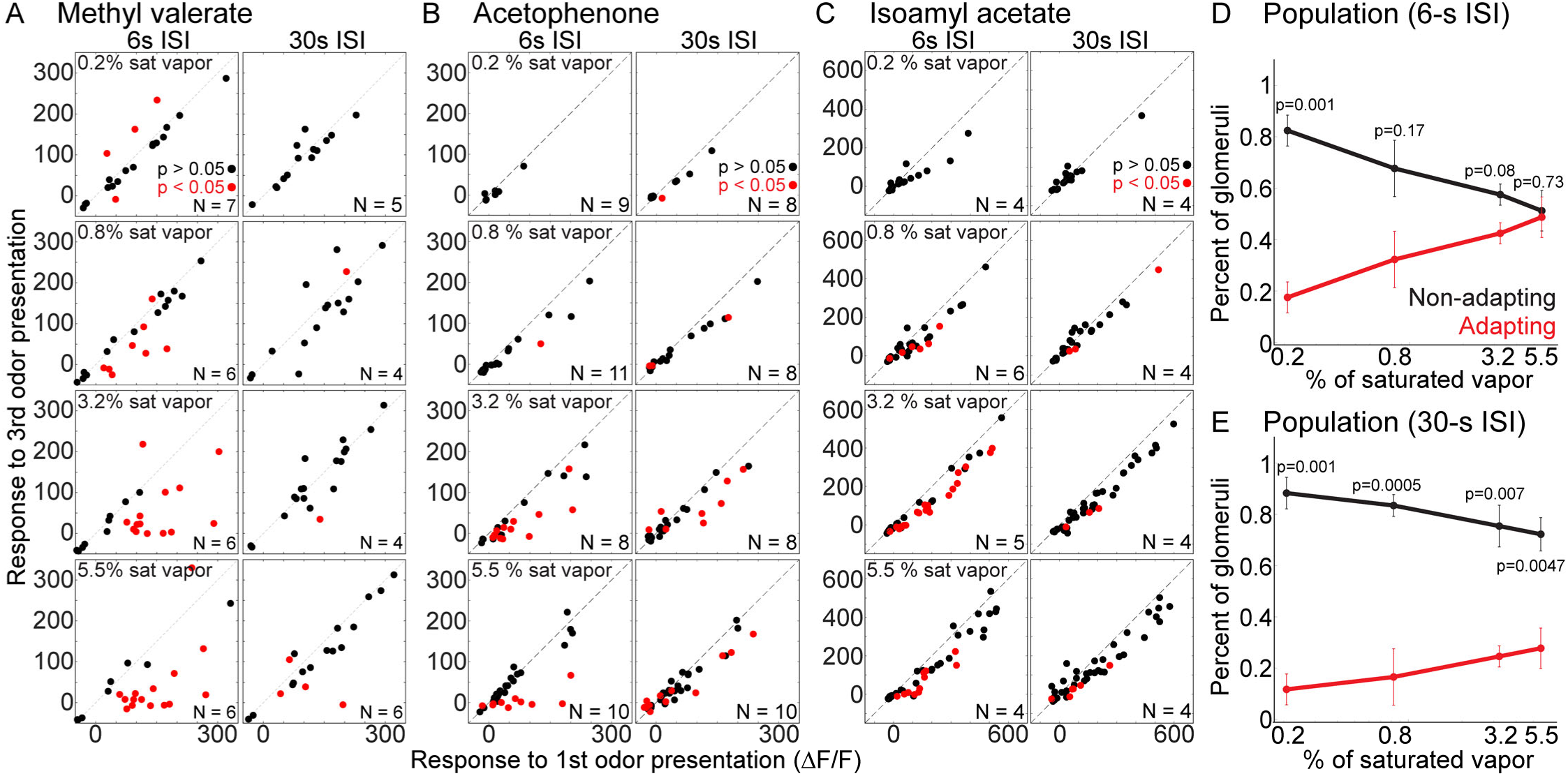
(A-C) Adaptation in individual mitral/tufted glomeruli in three preparations at different concentrations (rows). Each marker indicates the mean response of a single glomerulus. Red markers indicate glomeruli with a significantly different response to the 1st and 3rd odor presentation. The number of single trials used to determine statistical significance is in the bottom right of each panel. (D-E) The mean percent of responsive glomeruli in each preparation exhibiting significant adaptation (red lines) or stable responses (black lines) in response to the 1st to the 3rd odor presentation for 6-s (D) and 30-s (E) interstimulus intervals.

We examined the effect of repeated odor presentation across populations of glomeruli in 19 preparation-odor pairings in which 463 / 806 glomeruli exhibited a significant odor response at some concentration (42.4 ± 3.5 glomeruli were identified per preparation; between 21-64 per preparations, the analysis includes the preparations in **Fig. 9**). Each odor-concentration pairing was sampled in a similar number of trials for both 6-s and 30-s interstimulus interval conditions (6-s: 5.2 ± 0.35 trials; 30-s: 4.5 ± 0.38 trials per preparation; p > 0.31; range of 1-17 trials per condition).

The mean odor response from all glomeruli in this dataset, and the mean glomerular response for each preparation-odor pairing are illustrated in scatterplots (**Fig. 10A-B**). Consistent with the individual preparation examples in **Fig. 9**, lower concentrations and shorter interstimulus intervals evoked similar responses to each odor presentation, while higher concentrations increased the difference (**Fig. 10A-B**).

**Figure 10:**
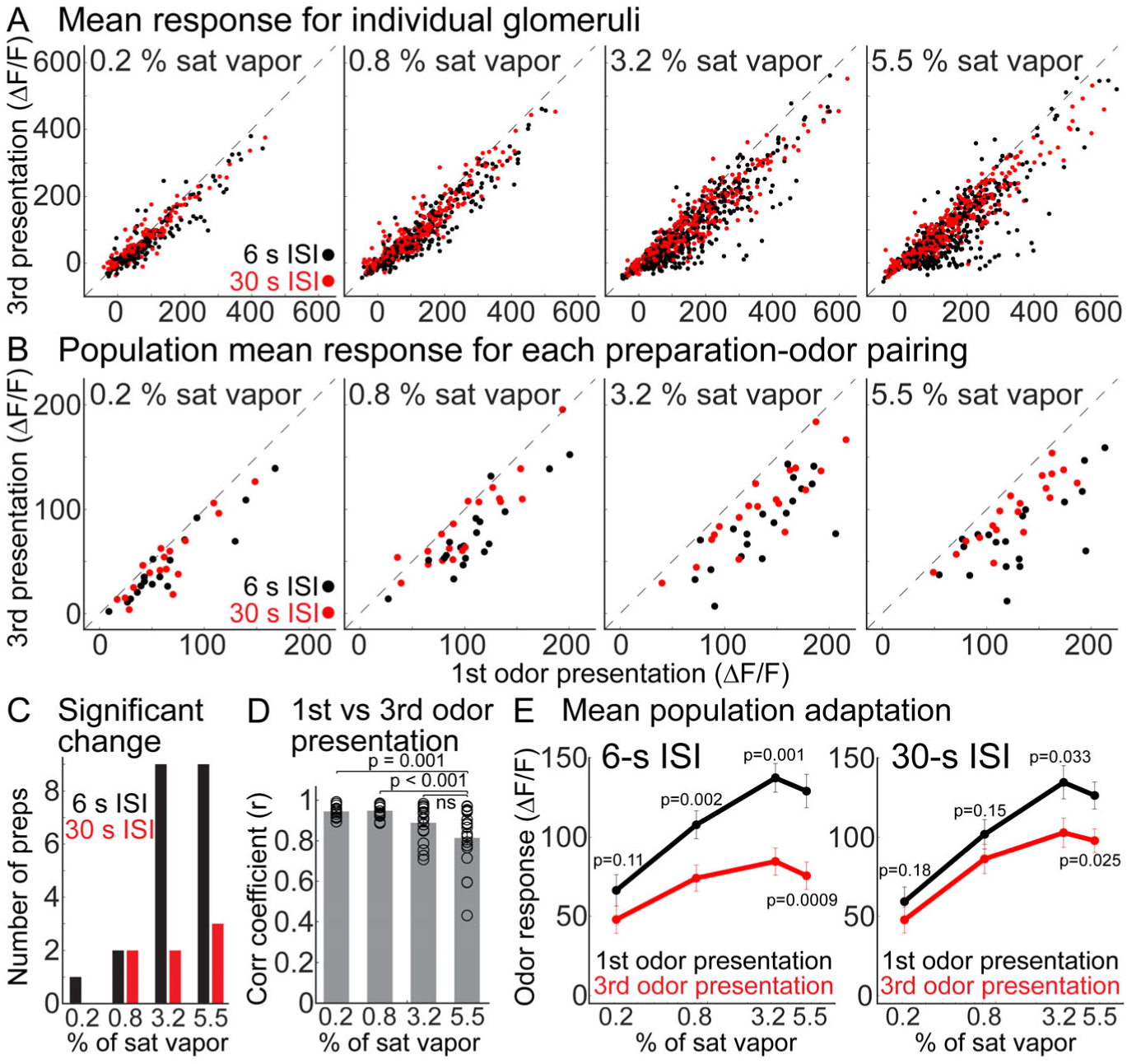
(A-B) Mean of all individual glomeruli (A) and preparation-odor pairings (B) in 19 preparation-odor pairings to the 1st and 3rd odor presentation at different concentrations. (C) Number of preparation-odor pairings from panel B in which the population response to the 3rd odor presentation was significantly different than the 1st. (D) Mean correlation of glomerular responses to the 1st and 3rd odor presentation. Each marker indicates a different preparation-odor pairing. (E) Mean response to the 1st and 3rd odor presentation as a function of odor concentration and interstimulus interval across all preparation-odor pairings.

The number of the individual preparation-odor pairings in which the mean glomerular response to the 3^rd^ odor presentation was significantly smaller than the 1^st^ presentation increased with higher concentrations for both interstimulus intervals (**Fig. 10C**). However, the 6-s interstimulus interval evoked significant adaptation in more preparations than the 30-s condition (**Fig. 10C**, black vs red bars). The population response to the 1^st^ and 3^rd^ odor presentation was highly correlated at lower concentrations and became increasingly decorrelated at higher concentrations (**Fig. 10D**, markers indicate the mean correlation for individual preparation-odor-concentration pairings, p < 0.01 for comparisons of 5.5% vs 0.8% and 5.5% and 0.2 % of saturated vapor). For the 6-s interstimulus interval, the mean response to the 3^rd^ odor presentation was significantly smaller in response to odors presented at 0.8%, 3.2% and 5.5% of saturated vapor (**Fig. 10E**, p-values are indicated in the panel). Extending the interstimulus interval to 30-s caused a partial recovery, although there was still a significant decrease at the two highest concentrations (**Fig. 10E**, p-values are indicated in the panel). The response amplitudes evoked by the 1^st^ odor presentation were not significantly different in these two groups of trials (**Fig. 10E**, comparison of black lines in 6-s and 30-s, p-values ranged between 0.35-0.96 for individual preparation comparisons).

We tested whether glomeruli suppressed by odors presented at 5.5% of saturated vapor have distinct adapting properties from excited glomeruli. For odors delivered with a 6-s interstimulus interval, suppressed glomeruli responded similarly to the 1^st^ and 3^rd^ odor presentation with response amplitudes of −21.7 ± 1.8 and −19.6 ± 2.2, respectively (p = 0.34, N = 41 glomeruli). However, the excited population includes polarity-switching glomeruli, which transition from excitation to suppression with repeated presentations (N = 53/463 excited glomeruli; 11.5% of responsive glomeruli at the highest concentration; 2.8 ± 0.8 per preparation; range of 0-11). The mean response of polarity switching glomeruli to the 1^st^ odor presentation was not significantly different from other excited glomeruli (response to 1^st^ presentation: polarity switching 149.7 ± 12.2; other excited 134 ± 6.1; p = 0.07). Both groups of glomeruli exhibited significant adaptation to the 3^rd^ odor presentation, although the magnitude of the polarity switching adaptation was ∼3-fold greater (response to 3^rd^ presentation: polarity switching 28.1 ± 6.4; other excited 87.4 ± 5.4; p < 0.00001 for both comparisons). Therefore, glomeruli that transition from excitation to suppression are a major source of adaptation in our data set, but do not exclusively mediate adaptation.

### Olfactory receptor neuron glomeruli exhibit minimal adaptation

In principle, mitral/tufted adaptation could reflect changes inherited from the olfactory receptor neuron input. To test this possibility, we carried out an analysis of adaptation in olfactory receptor neuron glomeruli that expressed GCaMP6s using awake *in vivo* 2-photon imaging in a separate cohort of transgenic mice. GCaMP6s was targeted to olfactory receptor neurons by mating the OMP-tTA and tetO-GCaMP6s transgenic lines. The offspring that expressed both genes were confirmed to express GCaMP6s in the olfactory nerve layer and glomerular layer (**Fig. 11A**) (Yu et al., 2004; Huang et al., 2022). We restricted adaptation measurements in olfactory receptor neuron glomeruli to odors presented with a 6-s interstimulus interval at the concentrations that evoked the strongest adaptation in mitral/tufted glomeruli. A frame subtraction analysis and time course responses from individual glomeruli demonstrate that the overall activity pattern and response amplitudes were comparable in response to repeated presentations (**Fig. 11B-D**). The response to the 1^st^ and 3^rd^ odor presentation was quantified for all individual glomeruli in the data set, as well as the overall population mean for each preparation-odor pairing at three different concentrations (**Fig. 11E-F**). Most individual glomeruli, and the mean from each preparation-odor pairing exhibited minimal adaptation (326 glomerulus-odor pairings in 12 preparation-odor pairings across 8 different animal preparations, includes responses to methyl valerate, isoamyl acetate and acetophenone) (**Fig. 11E-F**).

**Figure 11:**
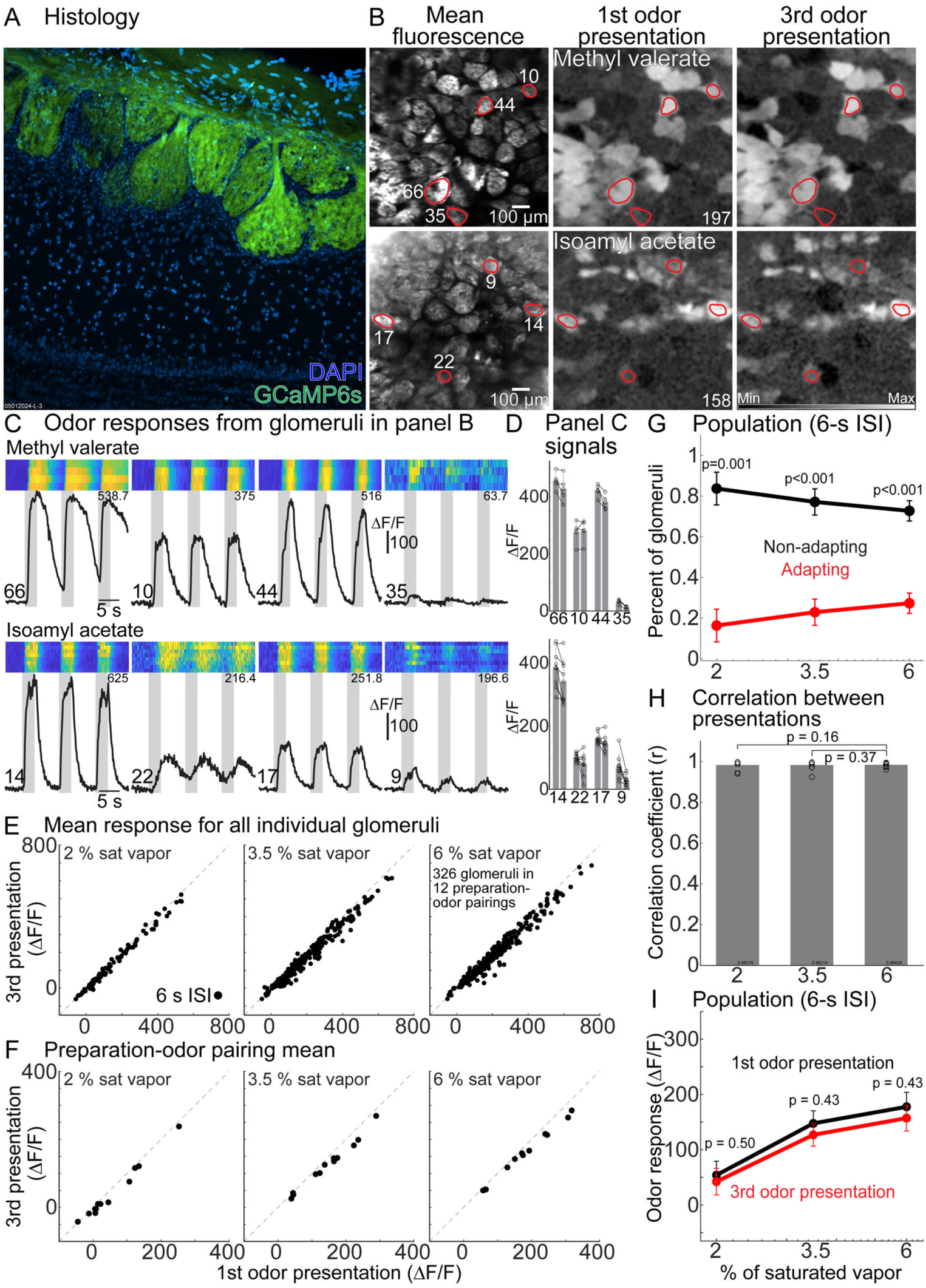
(A) Histology illustrating GCaMP6s expression in ORN glomeruli. (B) In vivo fluorescence and frame subtraction images to methyl valerate and isoamyl acetate (2% and 3.5% of sat vapor, respectively). The maximum ΔF/F value is indicated in the top right of each panel. (C) Single trial and mean odor responses from glomeruli in panel B. (D) Responses for the glomeruli in panel C. (E) Odor response from individual in 12 preparation-odor pairings. (F) Mean response from all glomeruli that responded to the 1st odor presentation for each preparation-odor pairing. (G) Percent of glomeruli in each preparation that exhibited significant adaptation (red lines) or responded similarly (black lines). (H) Correlation between odor presentations for the population response for each preparation-odor pairing. (I) Population mean across all preparation-odor pairings.

We similarly quantified the proportion of individual glomeruli that exhibited statistically significant adaptation in the same imaging field of view for preparations in which responses were measured in at least 4 single trials (includes 8 preparation-odor pairings; mean number of trials for the three tested concentrations: 8.4 ± 1.2, 8.6 ± 1.3 and 8.4 ± 1.3; 4-16 trials per condition). Although the proportion of olfactory receptor neuron glomeruli exhibiting significant adaptation increased at higher concentration, there were significantly more non-adapting glomeruli at all concentrations (**Fig. 11G**). The relationship of the response of individual glomeruli to the 1^st^ and 3^rd^ odor presentation was highly correlated (> 0.9) and did not change significantly as a function of concentration (**Fig. 11H**).

Although higher odor concentrations evoked significant adaptation in some olfactory receptor neuron glomeruli, the mean glomerular response to the 1^st^ and 3^rd^ odor presentation was not significantly different for individual preparation odor pairings (p-values ranged from 0.2 to 0.84). Consistently, the population mean response to the 1^st^ and 3^rd^ odor presentation was not statistically different for the three tested odor concentrations (**Fig. 11I**, p-values are indicated in the panel). Therefore, odors presented repeatedly with a 6-s interstimulus interval can evoke adaptation in individual olfactory receptor neuron glomeruli, although the population mean is not significantly impacted.

## Discussion

We used *in vivo* 2-photon Ca^2+^ imaging in the olfactory bulb of awake mice to measure how adaptation alters the responsiveness of mitral/tufted glomeruli. The same odor-concentration pairings evoked stable responses across different imaging trials separated by a minimum of 3 minutes, and across imaging sessions carried out on different days. Odors presented with shorter interstimulus intervals evoked adaptation non-uniformly across the glomerular population in a concentration-dependent manner. Within the same imaging field of view, some glomeruli adapted significantly while others did not. Higher concentrations increased the proportion of significantly adapting glomeruli, significantly decorrelated the glomerular population, and significantly attenuated the mean population response. Extending the interstimulus interval to 30-s reduced the proportion of adapting glomeruli and evoked less adaptation, yet the recovery across the glomerular population was incomplete.

Importantly, we demonstrate that the described adaptation is unlikely to reflect variations in respiration (**Fig. 5**), stimulus delivery (**Fig. 6**) or adaptation inherited from the periphery (**Fig. 11**). Therefore, recent odor exposure can impact the transmission of olfactory sensory processing from the bulb to the rest of the brain for upwards of 30 seconds. Adaptation on this timescale could be useful for mediating dynamic range adjustments to complex odor environments.

### Comparison of our results with previous studies

The result that mitral/tufted glomeruli exhibit relatively stable responses across different imaging sessions are in contrast with another study indicating that mitral/tufted glomeruli exhibit modest reductions in responsiveness over time (Kato et al., 2012). Although these differences could reflect methodological variations including a different transgenic mouse line (Pcdh21 versus Tbx21), optical sensor (GCaMP3 versus GCaMP6f), the frequency by which the odor was delivered, and the specific concentrations used, our results are consistent with other studies indicating that mitral/tufted glomeruli exhibit adaptation on shorter timescales and take upwards of 4 minutes to recover from recent odor stimulation (Ogg et al., 2015; Ogg et al., 2018; Storace and Cohen, 2021). Our results extend our previous report describing mitral/tufted glomerular adaptation to show that adaptation is present in awake mice, heterogeneous and concentration-dependent (Storace and Cohen, 2021). These data reconcile our work with previous results showing that individual mitral cells can exhibit stable responses since both adapting and non-adapting responses are present (Sobel and Tank, 1993; Wilson, 1998; Margrie et al., 2001; Kadohisa and Wilson, 2006).

However, the result that adaptation occurs heterogeneously is in contrast with previous work finding that adaptation occurs uniformly across the mitral/tufted glomerular population (Ogg et al., 2015). This difference may reflect the use of epifluorescence imaging where glomerular measurements can be influenced by out-of-focus fluorescence from neighboring glomeruli. In comparison, 2-photon imaging provides improved optical sectioning in the lateral and axial dimensions, minimizing the impact of out-of-focus signal. Indeed, efforts to correct for diffuse fluorescence in epifluorescence measurements by subtracting the background signal from each glomerulus yielded less uniform adaptation (Storace and Cohen, 2021).

Previous studies have reported adaptation in olfactory receptor neuron axon terminals, albeit either at much higher odor concentrations than used in the present study, or in response to high frequency sniffing (Wachowiak and Cohen, 2001; Verhagen et al., 2007; Carey et al., 2009; Lecoq et al., 2009). The mitral/tufted glomerular adaptation described here is unlikely to be explained by variations in respiration since adaptation often occurred over multiple inhalations and mice did not significantly change their respiratory patterns during our recording trials (**Fig. 5**). Although mice investigate novel odors with high-frequency sniffing, non-novel odors evoke respiratory rates that are comparable to our measurements (**Fig. 5**) (Verhagen et al., 2007; Wesson et al., 2008a; Wesson et al., 2008b). Furthermore, our findings of minimal adaptation in olfactory receptor neuron axon terminals and high correlation across odor repetitions are consistent with previous work from our laboratory and others (Chu et al., 2017; Storace and Cohen, 2021; Platisa et al., 2022).

### Methodological considerations

Measurements from olfactory receptor neuron and mitral/tufted glomeruli reflect a population average of the input from one olfactory receptor type and the apical dendrites of the mitral/tufted cells innervating that glomerulus. Glomerular measurements are therefore a useful complement to single cell recordings because it is difficult to determine whether individual mitral or tufted cells are connected to the same glomerulus, and different responses across nearby mitral/tufted cells could reflect different sensory input (Nagayama et al., 2007; Dhawale et al., 2010; Kikuta et al., 2013).

Because our study used the Tbx21-Cre transgenic line, our measurements will reflect an average of both mitral and tufted cells (Allen et al., 2007; Mitsui et al., 2011; Kosaka and Kosaka, 2012; Haddad et al., 2013; Koldaeva et al., 2021; Storace and Cohen, 2021), which are functionally distinct populations of projection neurons (Fukunaga et al., 2012; Igarashi et al., 2012). Future experiments incorporating transgenic animals with expression restricted to either cell population will be useful for better differentiating between the response properties of mitral versus tufted cells (Koldaeva et al., 2021). Action potentials generated in mitral cell apical dendrites propagate to the soma, and somatically generated action potentials backpropagate to the apical dendrites (Djurisic et al., 2004). However, subthreshold excitatory postsynaptic potentials can evoke small calcium increases, and TTX application incompletely blocks calcium signals measured from their apical dendrites in response to bath application of glutamate (Charpak et al., 2001; Kato et al., 2012).

Therefore, calcium measurements from the apical dendrites of mitral/tufted cells innervating a glomerulus includes both pre- and postsynaptic responses. Future experiments would benefit from development of genetically encoded voltage indicators that are tuned to supra-threshold voltage changes to allow for unambiguous optical measurements of spiking activity from mitral/tufted apical dendrites innervating a glomerulus (Leong and Storace, 2024).

Our PID analysis illustrates that increasing the air flow speed through the odor vial increased the PID signal (**Fig. 6**). Notably, a ∼100-fold dilution of methyl valerate in mineral oil presented at 10 % of saturated reduced the PID signal to a level comparable to what was measured in response to the pure odor delivered at 3 % of saturated vapor. Although additional studies are needed that compare the effect of how liquid dilutions alter the PID signal for different odors, our result suggests that comparing concentration measurements across studies requires understanding how the odor was prepared, as well as the absolute flow rates that pass through the odor vial (Jennings et al., 2023).

In this study we compared adapting responses in glomeruli measured from mitral/tufted cells and olfactory receptor neurons using the optical sensors GCaMP6f and GCaMP6s, respectively. Each calcium indicator has distinct biophysical properties that will contribute to measured differences (Akerboom et al., 2012; Sun et al., 2013; Badura et al., 2014; Storace et al., 2015). However, the present data are consistent with our previous study that reported that olfactory receptor neurons exhibit less adaptation than mitral/tufted glomeruli measured using an organic calcium dye, and GCaMP3 and GCaMP6f (Storace and Cohen, 2021).

### Mechanisms underlying adaptation and functional relevance

The presence of glomeruli with qualitatively distinct adapting properties suggests that multiple mechanisms may be involved in mitral/tufted glomerular adaptation. Recurrent inhibition can shape the spiking activity of mitral cells in a manner that depends on the overall activation of the mitral cell itself (Margrie et al., 2001). The mechanism is likely to be complex since granule cells exhibit strong paired pulse depression, suggesting that strongly activated granule cells should provide progressively weaker inhibition onto their synaptic contacts (Dietz and Murthy, 2005). The specific combination of receptors expressed on different cell types is also likely to be important since NMDA antagonism can impair habituation to odor stimuli *in vivo* (Chaudhury et al., 2010). Feedback from other brain areas is another candidate mechanism since stimulating acetylcholine input to the bulb dishabituates glomerular and behavioral sensitivity (Ogg et al., 2018). The presence of a concentration-dependent form of adaptation is compatible with a model of lateral inhibition whereby higher concentrations of an odor must drive input to increasing numbers of glomeruli by activating lower affinity olfactory receptors. This process should increase the magnitude of lateral inhibition in the bulb (Wachowiak and Cohen, 2001; Banerjee et al., 2015; Storace and Cohen, 2017; Storace et al., 2019). Such a mechanism could account for the heterogeneity of different adapting responses since each glomerulus is likely to be differently influenced by lateral circuits (Fantana et al., 2008). This hypothesis can be tested in future studies by modulating the magnitude of lateral inhibition within the circuit by selectively manipulating specific olfactory receptor neuron or interneuron types within the bulb (Banerjee et al., 2015; Braubach et al., 2018).

Our result that glomeruli suppressed to the 1^st^ odor presentation exhibit minimal adaptation raises the possibility that the mechanisms underlying suppression and adaptation are distinct. However, suppressed calcium signals in mitral/tufted glomeruli likely reflect a decrease in spontaneous activity in response to the odor stimulus. Consequently, there may be a floor effect in which additional adaptive processes cannot be detected. Indeed, a notable subpopulation of excited glomeruli transitioned from excitation to suppression. Future studies measuring suppressed responses at higher concentrations, and with a larger number of polarity transitioning glomeruli are required to test whether adaptation uniquely impacts suppressed glomeruli.

### Estimating variability within the mitral/tufted glomerular population

Neural measurements in awake mice are more variable than in anesthetized preparations in part due to the variable nature of respiration in awake mice. Although it is possible to control the frequency of respiration in anesthetized preparations, such methods are challenging to implement in awake preparations (Short and Wachowiak, 2019; Eiting and Wachowiak, 2020). Here we purposely oversampled the number of trials for each odor-concentration pairing, finding that 4 subsamples are needed to recapitulate a reasonable estimate of the response from a glomerulus measured in a larger number of samples. This result may provide a useful starting point for establishing sample sizes for future experiments.

### Conclusions

Here we extend previous work describing a form of adaptation in mitral/tufted glomeruli that occurs on timescales that could be relevant for processes related to odor-background segmentation (Gottfried, 2010). However, future studies are needed to confirm the functional and behavioral relevance of adaptation in mitral/tufted glomeruli. Simultaneous comparisons of olfactory receptor neuron input and mitral/tufted glomerular output will provide insight into the nature of how the olfactory bulb can transform a stable sensory input into an adapting output. Comparisons of mitral/tufted glomeruli before and after adaptation has taken place will define precisely how adaptation shapes future responsiveness of the olfactory bulb circuit (Parabucki et al., 2019; Benda, 2021; Martelli and Storace, 2021). Finally, although there is evidence that mice behaviorally habituate to odors presented on similar timescales, simultaneous imaging from mitral/tufted glomeruli during a behavioral assay is necessary to clearly link glomerular measurements with perception.

## Methods

### Transgenic mice

GCaMP6f was targeted to mitral/tufted glomeruli by mating the Ai148 GCaMP6f transgenic reporter line (Jax stock #030328) to the Tbx21-cre transgenic line (Jax stock #024507). GCaMP6s was targeted to olfactory receptor neuron glomeruli by mating the tetO-GCaMP6s transgenic reporter line (Jax stock #024742) to the OMP-tTA transgenic line (Jac stock #017754) (Huang et al., 2022).

Offspring that expressed eGFP and Cre recombinase (mitral/tufted glomeruli), or eGFP and tTA (olfactory receptor neurons glomeruli) were used for experiments. Genotyping was performed by Transnetyx (Cordova, TN). Appropriate targeting of either GCaMP to olfactory receptor neurons or mitral/tufted glomeruli was confirmed histologically in a subset of the preparations based on endogenous fluorescence expression.

### Surgical procedures

All procedures were approved by the Florida State University Animal Care and Use Committee. Male and female adult (> 21 days) transgenic mice were anesthetized using ketamine/xylazine (90 / 10 mg/kg, Zoetis, Kalamazoo, MI), placed on a heating pad and had ophthalmic ointment applied to their eyes. Mice were given a pre-operative dose of carprofen (10 mg/kg, Zoetis, Kalamazoo, MI), atropine (0.2 mg/kg, Covetrus, Dublin, OH), dexamethasone (4 mg/kg, Bimeda, La Sueur, MN), and bupivacaine (2.5 mg/kg, Hospira, Lake Forest, IL). Fur was removed using a depilatory agent, and the skin was scrubbed with 70% isopropyl alcohol and iodine (Covidien, Mansfield, MA). An incision was made to remove the skin over the skull and blunt dissection was used to remove the underlying membrane. Dental cement (Metabond, Covetrus, Dublin, OH) was used to attach a custom headpost to the skull, which was held using a headpost holder. The bone above the olfactory bulb was either thinned using a dental drill (Osada, XL-230, Los Angeles, CA) and covered with cyanoacrylate to improve optical clarity or was removed and replaced with #1 cover glass. Upon completion of the surgery, the mouse was allowed to recover on a heating pad until they were ambulatory. Animals were given additional analgesic at the end of the day of surgery and for at least 3 days post-operatively.

### Histology

Mice were euthanized (euthasol) following imaging and either underwent cardiac perfusion with phosphate buffered saline and 4% paraformaldehyde or had their brains extracted and post-fixed in 4% paraformaldehyde before being cut on a vibratome in 40 µm sections (Leica VT1000S, Deer Park, IL). Sections through the olfactory bulb were mounted on slides and were coverslipped using Fluoromount-G containing DAPI (SouthernBiotech, Birmingham, AL). Endogenous fluorescence expression of GCaMP6f or GCaMP6s was observed using a GFP filter set on either a Zeiss Axioskop epifluorescence microscope or a Nikon CSU-W1 spinning disk confocal microscope.

### 2-photon imaging

2-photon imaging was performed using a Sutter MOM 2-photon microscope equipped with an 8 kHz (30.9 Hz) resonant scanner (Cambridge Technology, USA) and an emission pathway equipped with a GaAsP PMT (#H10770PA-40-04, Hamamatsu, Japan). Laser excitation was provided using a Spectra-Physics DS+ between 940-980 nm with power modulated by a Pockels cell (Model #350-105-02, Conoptics, Danbury, CT), or an Alcor 920 (920 nm) with power controlled by an internal acousto-optic modulator. Imaging was performed using a Nikon 16x 0.8 N.A. or the Cousa 10x 0.5 N.A. objective lens (Yu et al., 2024). Laser power was confirmed to be less than 150 mW at the output of the objective lens measured using a power meter (Newport 843-R) for scanning areas ranging between 711 µm^2^ (16 x lens) and 1138 µm^2^ (10x lens).

### Odorant delivery

Odorants included: methyl valerate (CAS #624-24-8), isoamyl acetate (CAS #123-92-2), benzaldehyde (100-52-7), ethyl butyrate (CAS #105-54-4), acetophenone (CAS #100-52-7), and 2-phenylethanol (CAS #60-12-8) (Sigma-Aldrich, USA) were used at concentrations between 0.05 and 6 % of saturated vapor. For mitral/tufted glomerular measurements, the olfactometer design involved air being pushed through vials of pure odor using a syringe pump (NE-1000, PumpSystems, Farmingdale, NY) running at different flow rates (0.25 – 28 ml /min). This odor stream underwent an initial air dilution with a lower flow rate of clean air (30 ml/min). The resulting odorized air stream connected to a dual 3-way solenoid valve (360T041, NResearch, West Caldwell, NJ), which was connected to an exhaust, a clean air stream, and a delivery manifold which served as the final delivery apparatus. The delivery valve was connected to a Teflon delivery manifold placed in front of the mouse’s nose, which had a higher flow rate of clean air constantly flowing through it (450 ml/min). Prior to odor triggering, the solenoid sent the odorized air stream to the exhaust, and the clean air into the delivery manifold, while triggering it caused the odor to be injected into the delivery manifold where it underwent a second air dilution. For olfactory receptor neuron glomerular measurements, the olfactometer design was similar except air flow was controlled by mass flow controllers (Alicat, MC-100SCCM and MC-1SLPM), and underwent only a single air dilution step (Williams and Dewan, 2020).

The odor delivery time-course for both odor delivery systems were confirmed using a photoionization detector set to 1x gain, and pump speed set to high (PID, 200C, Aurora Scientific, Aurora, ON). Three different odor vials containing either 10 ml of pure methyl valerate, a 1:10 dilution of methyl valerate (1 ml odor + 9 ml mineral oil), or a 1:100 dilution (0.1 ml odor + 9.9 ml mineral oil). The PID signals in response to different air and liquid dilutions were recorded directly from the PID into the Sutter MScan system. All traces had a 0.18 mV DC component which was subtracted in MATLAB to reference the PID baseline to zero. PID amplitudes were calculated as the average voltage during a 2400 msec window during the odor stimulus.

### Imaging procedures

Prior to data collection mice were positioned underneath the microscope and the angle of the headpost holder was adjusted to optimize the imaging field of view. The headpost holder position was then locked into place so that the mouse could be precisely realigned during future imaging sessions. Only one field of view was imaged from each mouse preparation. During data collection, awake head-fixed mice were placed underneath the microscope objective with the olfactometer and a thermocouple (to measure respiration, Omega 5TC-TT-K-36-36, Newark) near its nose. The signals from the respiration sensor were amplified and low-pass filtered using a differential amplifier (Model 3000, AM-Systems, Sequim, WA), which was simultaneously recorded by the imaging system at 1000 Hz. Different odor-concentration pairings were presented with interstimulus intervals of 6, 12 and 30 seconds. Individual trials were separated by a minimum of 3 minutes. For mitral/tufted glomerular recordings, odor-concentration pairings included 4 steps between 0.2 – 5.5% of saturated vapor.

Measuring the response to the full range of concentrations for a particular odor was prioritized within a single imaging session for at least one interstimulus interval. The response to the same odor-concentration-interstimulus interval pairing was typically measured in at least two consecutive trials to assess within-day, across-trial repeatability. If time permitted, the response to the same odor-concentration pairings were measured to a different interstimulus interval.

### Data analysis

Following data acquisition, the raw image files were spatially and temporally averaged from 512×512 pixels sampled at 30.9 Hz to 256×256 pixels sampled at 7.72 Hz. The resulting data were exported to TIFF format for all subsequent analysis. Because the mice were typically calm, motion correction was not used. However, occasional recordings with sufficient motion artifact that made it impossible to interpret the measurements were discarded from the subsequent analysis pipeline.

Segmentation of image stacks into glomerular regions of interest was manually carried out in custom software (Turbo-SM, SciMeasure, Decatur, GA). Glomeruli were identified as glomerular sized regions of interest using the mean fluorescence, as well as a frame subtraction analysis that displays the difference between the frames during and prior to the odor stimulation. Region of interest overlays were typically identified on the first day, and if necessary, were adjusted to account for minor changes in positioning across trials and imaging sessions. The pixel areas containing the regions of interest were saved and the fluorescence time course values from each region of interest were extracted for subsequent analysis. All fluorescence time course traces were converted to ΔF/F by dividing each trace by the mean of all the frames prior to the odor command trigger (typically 23).

The mean fluorescence and frame subtraction images are from the average of at least two single trials (e.g., **Fig. 1B**, **Fig. 3B-D**, **Fig. 11B**). The mean fluorescence images are generated from the average of all the frames during the imaging trial. The frame subtraction images were generated by subtracting the average of 19 frames during odor stimulation from the average of the 9 frames prior to the stimulus (**Fig. 1B**, *bottom row*; **Fig. 3B-D**, *1^st^ and 3^rd^ presentation*). The frame subtraction difference images were generated by subtracting the average of the 20 frames during the 3^rd^ odor presentation from the average of the 20 frames during the 1^st^ odor presentation (**Fig. 3B-D**, *difference map*). The frame subtraction images underwent two passes of a low-pass spatial filter and were converted to ΔF/F by dividing the fluorescence value of each pixel by the mean of at least 60 consecutive frames in the image stack in Turbo-SM. The colorscale minimum to maximum range is fixed for the two different odors in **Fig. 1B** and the individual range of values for each image is indicated at the bottom of each panel.

Odor responses (ΔF/F) were calculated as the largest difference between a 1200 msec temporal window during the peak of the odor response and the time prior to odor stimulation. This value was calculated independently for each glomerulus (**Fig. 4B, D, F**; **Fig. 7B**, **D**; **Fig. 11D**). All statistical comparisons were performed using the Wilcoxon rank sum test (ranksum function in MATLAB), and the Kruskal-wallis test (kruskalwallis function in MATLAB).

Across trial correlations were calculated by measuring the correlation between the 2.5 seconds prior to odor stimulation, and during odor stimulation for individual traces (**Fig. 1E**), and for the mean population response across all glomeruli in a field of view (**Fig. 1F**). The mean glomerular output correlation analysis (**Fig. 2C**) was generated by averaging the correlation coefficients from all trials on the same day or all trial pairings that took place on different days.

For comparisons of amplitude across days, glomeruli were selected for analysis if they exhibited a minimum of a 3 standard deviation change to odor stimulation across all imaging sessions (thresholds ranged between 3-6, 4.6 ± 0.3, N = 16 preparation-odor pairings). Glomeruli that did not appear similarly in the baseline fluorescence (mean of the first 23 frames) on all imaging days were excluded from the analysis (Kato et al., 2012). For the glomeruli that met these criteria, the mean response to 3.2 % of saturated vapor was calculated for each imaging session for each preparation-odor pairing (**Fig. 1D**, **Fig. 2A-B**).

Because the number of imaging sessions, and the time between each imaging session was not the same for each preparation, population responses are illustrated binned across different imaging sessions or based on the number of days since the 1^st^ imaging session (**Fig. 2A-B**, the number of preparation-odor pairings included in each bin are indicated at the bottom of each graph).

The subsampling analysis was performed by generating 1000 random subsamples of trials from all trials for each preparation-odor pairing at 3.2% of saturated vapor (datasample function with replacement in MATLAB). The proportion of the subsamples whose mean was within 1 standard deviation of the unsampled mean was calculated for each glomerulus. This proportion was averaged across all glomeruli to determine the mean proportion across the glomerular population. This process was repeated for different subsample sizes from 1-5. Increasing the numbers of random subsamples yielded similar results.

For the respiration analysis in **Fig. 5**, individual inhalations were identified in the respiration traces using the islocalmax function in MATLAB. Inhalation counts were measured by counting the number of inspirations while the odor command was on. Inter-inhalation intervals were calculated by measuring the time between subsequent inhalations using the diff function in MATLAB.

The adaptation analysis was performed on glomeruli if they responded to the 1^st^ odor presentation with at least a 3 standard deviation change from the baseline fluorescence. The mean number of significantly adapting and non-adapting glomeruli was calculated based on the number of glomeruli exhibiting a statistically significant change to the odor stimulation (**Fig. 9D-E**, left panel).

Preparations were only included in this analysis if measurements were sampled in a minimum of 4 trials for all odor-concentration conditions (**Fig. 9D-E**). The percentage of significantly adapting glomeruli was calculated by dividing the number of glomeruli exhibiting a statistically significant change by the sum of all responsive glomeruli (**Fig. 9D-E**).

## Acknowledgements

This work was supported by funding from NIDCD R01 DC020519, and internal funds from Florida State University. We are grateful to Dr. Evan Lloyd for critical comments on an early draft of this manuscript.

## Data Sharing and Data Availability

Data will be made available upon request from the corresponding author.

## References

Akerboom J et al. (2012) Optimization of a GCaMP calcium indicator for neural activity imaging. J Neurosci 32:13819–13840.

Allen ZJ, Waclaw RR, Colbert MC, Campbell K (2007) Molecular identity of olfactory bulb interneurons: transcriptional codes of periglomerular neuron subtypes. J Mol Histol 38:517–525.

Badura A, Sun XR, Giovannucci A, Lynch LA, Wang SS (2014) Fast calcium sensor proteins for monitoring neural activity. Neurophotonics 1:025008.

Banerjee A, Marbach F, Anselmi F, Koh MS, Davis MB, Garcia da Silva P, Delevich K, Oyibo HK, Gupta P, Li B, Albeanu DF (2015) An Interglomerular Circuit Gates Glomerular Output and Implements Gain Control in the Mouse Olfactory Bulb. Neuron 87:193–207.

Benda J (2021) Neural adaptation. Curr Biol 31:R110–R116.

Braubach O, Tombaz T, Geiller T, Homma R, Bozza T, Cohen LB, Choi Y (2018) Sparsened neuronal activity in an optogenetically activated olfactory glomerulus. Sci Rep 8:14955.

Carey RM, Verhagen JV, Wesson DW, Pirez N, Wachowiak M (2009) Temporal structure of receptor neuron input to the olfactory bulb imaged in behaving rats. J Neurophysiol 101:1073–1088.

Charpak S, Mertz J, Beaurepaire E, Moreaux L, Delaney K (2001) Odor-evoked calcium signals in dendrites of rat mitral cells. Proc Natl Acad Sci U S A 98:1230–1234.

Chaudhury D, Manella L, Arellanos A, Escanilla O, Cleland TA, Linster C (2010) Olfactory bulb habituation to odor stimuli. Behav Neurosci 124:490–499.

Chu MW, Li WL, Komiyama T (2017) Lack of Pattern Separation in Sensory Inputs to the Olfactory Bulb during Perceptual Learning. eNeuro 4.

Daigle TL et al. (2018) A Suite of Transgenic Driver and Reporter Mouse Lines with Enhanced Brain-Cell-Type Targeting and Functionality. Cell 174:465–480 e422.

Dhawale AK, Hagiwara A, Bhalla US, Murthy VN, Albeanu DF (2010) Non-redundant odor coding by sister mitral cells revealed by light addressable glomeruli in the mouse. Nat Neurosci 13:1404–1412.

Dietz SB, Murthy VN (2005) Contrasting short-term plasticity at two sides of the mitral-granule reciprocal synapse in the mammalian olfactory bulb. J Physiol 569:475–488.

Djurisic M, Antic S, Chen WR, Zecevic D (2004) Voltage imaging from dendrites of mitral cells: EPSP attenuation and spike trigger zones. J Neurosci 24:6703–6714.

Duchamp-Viret P, Chaput MA, Duchamp A (1999) Odor response properties of rat olfactory receptor neurons. Science 284:2171–2174.

Eiting TP, Wachowiak M (2020) Differential Impacts of Repeated Sampling on Odor Representations by Genetically-Defined Mitral and Tufted Cell Subpopulations in the Mouse Olfactory Bulb. J Neurosci 40:6177–6188.

Fantana AL, Soucy ER, Meister M (2008) Rat olfactory bulb mitral cells receive sparse glomerular inputs. Neuron 59:802–814.

Fried HU, Fuss SH, Korsching SI (2002) Selective imaging of presynaptic activity in the mouse olfactory bulb shows concentration and structure dependence of odor responses in identified glomeruli. Proc Natl Acad Sci U S A 99:3222–3227.

Fukunaga I, Berning M, Kollo M, Schmaltz A, Schaefer AT (2012) Two distinct channels of olfactory bulb output. Neuron 75:320–329.

Gottfried JA (2010) Central mechanisms of odour object perception. Nat Rev Neurosci 11:628–641.

Gross-Isseroff R, Lancet D (1988) Concentration-dependent changes of perceived odor quality. Chemical Senses 13:191-204.

Haddad R, Lanjuin A, Madisen L, Zeng H, Murthy VN, Uchida N (2013) Olfactory cortical neurons read out a relative time code in the olfactory bulb. Nat Neurosci 16:949–957.

Homma R, Cohen LB, Kosmidis EK, Youngentob SL (2009) Perceptual stability during dramatic changes in olfactory bulb activation maps and dramatic declines in activation amplitudes. Eur J Neurosci 29:1027–1034.

Huang JS, Kunkhyen T, Rangel AN, Brechbill TR, Gregory JD, Winson-Bushby ED, Liu B, Avon JT, Muggleton RJ, Cheetham CEJ (2022) Immature olfactory sensory neurons provide behaviourally relevant sensory input to the olfactory bulb. Nat Commun 13:6194.

Igarashi KM, Ieki N, An M, Yamaguchi Y, Nagayama S, Kobayakawa K, Kobayakawa R, Tanifuji M, Sakano H, Chen WR, Mori K (2012) Parallel mitral and tufted cell pathways route distinct odor information to different targets in the olfactory cortex. J Neurosci 32:7970–7985.

Jennings L, Williams E, Caton S, Avlas M, Dewan A (2023) Estimating the relationship between liquid-and vapor-phase odorant concentrations using a photoionization detector (PID)-based approach. Chem Senses 48.

Kadohisa M, Wilson DA (2006) Olfactory cortical adaptation facilitates detection of odors against background. J Neurophysiol 95:1888–1896.

Kato HK, Chu MW, Isaacson JS, Komiyama T (2012) Dynamic sensory representations in the olfactory bulb: modulation by wakefulness and experience. Neuron 76:962–975.

Kikuta S, Fletcher ML, Homma R, Yamasoba T, Nagayama S (2013) Odorant response properties of individual neurons in an olfactory glomerular module. Neuron 77:1122–1135.

Koldaeva A, Zhang C, Huang YP, Reinert JK, Mizuno S, Sugiyama F, Takahashi S, Soliman T, Matsunami H, Fukunaga I (2021) Generation and Characterization of a Cell Type-Specific, Inducible Cre-Driver Line to Study Olfactory Processing. J Neurosci 41:6449–6467.

Kosaka T, Kosaka K (2012) Further characterization of the juxtaglomerular neurons in the mouse main olfactory bulb by transcription factors, Sp8 and Tbx21. Neurosci Res 73:24-31.

Kurahashi T, Menini A (1997) Mechanism of odorant adaptation in the olfactory receptor cell. Nature 385:725–729.

Lecoq J, Tiret P, Charpak S (2009) Peripheral adaptation codes for high odor concentration in glomeruli. J Neurosci 29:3067–3072.

Leong LM, Storace DA (2024) Imaging different cell populations in the mouse olfactory bulb using the genetically encoded voltage indicator ArcLight. Neurophotonics 11:033402.

Margrie TW, Sakmann B, Urban NN (2001) Action potential propagation in mitral cell lateral dendrites is decremental and controls recurrent and lateral inhibition in the mammalian olfactory bulb. Proc Natl Acad Sci U S A 98:319–324.

Martelli C, Storace DA (2021) Stimulus Driven Functional Transformations in the Early Olfactory System. Frontiers in Cellular Neuroscience 15.

Mitsui S, Igarashi KM, Mori K, Yoshihara Y (2011) Genetic visualization of the secondary olfactory pathway in Tbx21 transgenic mice. Neural Syst Circuits 1:5.

Mombaerts P, Wang F, Dulac C, Chao SK, Nemes A, Mendelsohn M, Edmondson J, Axel R (1996) Visualizing an olfactory sensory map. Cell 87:675–686.

Nagayama S, Zeng S, Xiong W, Fletcher ML, Masurkar AV, Davis DJ, Pieribone VA, Chen WR (2007) In vivo simultaneous tracing and Ca(2+) imaging of local neuronal circuits. Neuron 53:789–803.

Ogg MC, Bendahamane M, Fletcher ML (2015) Habituation of glomerular responses in the olfactory bulb following prolonged odor stimulation reflects reduced peripheral input. Front Mol Neurosci 8:53.

Ogg MC, Ross JM, Bendahmane M, Fletcher ML (2018) Olfactory bulb acetylcholine release dishabituates odor responses and reinstates odor investigation. Nat Commun 9:1868.

Parabucki A, Bizer A, Morris G, Munoz AE, Bala ADS, Smear M, Shusterman R (2019) Odor Concentration Change Coding in the Olfactory Bulb. eNeuro 6.

Platisa J, Zeng H, Madisen L, Cohen LB, Pieribone VA, Storace DA (2022) Voltage imaging in the olfactory bulb using transgenic mouse lines expressing the genetically encoded voltage indicator ArcLight. Sci Rep 12:1875.

Potter SM, Zheng C, Koos DS, Feinstein P, Fraser SE, Mombaerts P (2001) Structure and emergence of specific olfactory glomeruli in the mouse. J Neurosci 21:9713–9723.

Rinberg D, Koulakov A, Gelperin A (2006) Sparse odor coding in awake behaving mice. J Neurosci 26:8857–8865.

Short SM, Wachowiak M (2019) Temporal Dynamics of Inhalation-Linked Activity across Defined Subpopulations of Mouse Olfactory Bulb Neurons Imaged In Vivo. eNeuro 6.

Sobel EC, Tank DW (1993) Timing of odor stimulation does not alter patterning of olfactory bulb unit activity in freely breathing rats. J Neurophysiol 69:1331–1337.

Storace DA, Cohen LB (2017) Measuring the olfactory bulb input-output transformation reveals a contribution to the perception of odorant concentration invariance. Nat Commun 8:81.

Storace DA, Cohen LB (2021) The mammalian olfactory bulb contributes to the adaptation of odor responses: a second perceptual computation carried out by the bulb. eNeuro.

Storace DA, Cohen LB, Choi Y (2019) Using Genetically Encoded Voltage Indicators (GEVIs) to Study the Input-Output Transformation of the Mammalian Olfactory Bulb. Front Cell Neurosci 13:342.

Storace DA, Braubach OR, Jin L, Cohen LB, Sung U (2015) Monitoring brain activity with protein voltage and calcium sensors. Sci Rep 5:10212.

Sun XR, Badura A, Pacheco DA, Lynch LA, Schneider ER, Taylor MP, Hogue IB, Enquist LW, Murthy M, Wang SS (2013) Fast GCaMPs for improved tracking of neuronal activity. Nat Commun 4:2170.

Torre V, Ashmore JF, Lamb TD, Menini A (1995) Transduction and adaptation in sensory receptor cells. J Neurosci 15:7757–7768.

Uchida N, Mainen ZF (2007) Odor concentration invariance by chemical ratio coding. Front Syst Neurosci 1:3.

Verhagen JV, Wesson DW, Netoff TI, White JA, Wachowiak M (2007) Sniffing controls an adaptive filter of sensory input to the olfactory bulb. Nat Neurosci 10:631–639.

Wachowiak M, Cohen LB (2001) Representation of odorants by receptor neuron input to the mouse olfactory bulb. Neuron 32:723–735.

Wachowiak M, Economo MN, Diaz-Quesada M, Brunert D, Wesson DW, White JA, Rothermel M (2013) Optical dissection of odor information processing in vivo using GCaMPs expressed in specified cell types of the olfactory bulb. J Neurosci 33:5285–5300.

Wark B, Lundstrom BN, Fairhall A (2007) Sensory adaptation. Curr Opin Neurobiol 17:423–429.

Weber AI, Fairhall AL (2019) The role of adaptation in neural coding. Curr Opin Neurobiol 58:135-140.

Wesson DW, Carey RM, Verhagen JV, Wachowiak M (2008a) Rapid encoding and perception of novel odors in the rat. PLoS Biol 6:e82.

Wesson DW, Donahou TN, Johnson MO, Wachowiak M (2008b) Sniffing behavior of mice during performance in odor-guided tasks. Chem Senses 33:581–596.

Whitmire CJ, Stanley GB (2016) Rapid Sensory Adaptation Redux: A Circuit Perspective. Neuron 92:298–315.

Williams E, Dewan A (2020) Olfactory Detection Thresholds for Primary Aliphatic Alcohols in Mice. Chem Senses 45:513-521.

Wilson DA (1998) Habituation of odor responses in the rat anterior piriform cortex. J Neurophysiol 79:1425–1440.

Yu CH et al. (2024) The Cousa objective: a long-working distance air objective for multiphoton imaging in vivo. Nat Methods 21:132–141.

Yu CR, Power J, Barnea G, O’Donnell S, Brown HE, Osborne J, Axel R, Gogos JA (2004) Spontaneous neural activity is required for the establishment and maintenance of the olfactory sensory map. Neuron 42:553–566.

Zak JD, Reddy G, Vergassola M, Murthy VN (2020) Antagonistic odor interactions in olfactory sensory neurons are widespread in freely breathing mice. Nat Commun 11:3350.

